# A New Soaking Protocol to Live Stain *C. elegans* with Proteostat Dye Reveals Global Patterns of Stress-Induced Protein Aggregation

**DOI:** 10.64898/2026.09.30.755777

**Authors:** Tarah Krishingner, Sarah R. Fausett

**Author notes:** Department of Biology and Marine Biology, University of North Carolina Wilmington, Wilmington, NC, USA. Corresponding author (SF).

## Abstract

Protein aggregates are a diverse array of intra- and extracellular structures containing misfolded proteins with limited macromolecular movement. Protein aggregation is induced by aging and cellular stress and is a hallmark of neurodegenerative diseases including Huntington’s, Alzheimer’s, and Parkinson’s. Fluorescent tools to visualize global protein aggregation in whole organisms are, therefore, important for screening genes and compounds that may reduce or alter protein aggregation in connection with improved health outcomes. Fluorescent probes that report low fluidity structures in cells, such as molecular rotor dyes, are gaining popularity and overcome certain limitations of targeted markers such as tagged proteins or antibodies. However, permeability has been a key barrier preventing widespread use of such probes in whole organisms, including in *C. elegans*. To achieve visualization of non-specific protein aggregates in whole *C. elegans* nematodes without the need for strain engineering, we have developed a simple soaking protocol. Our protocol overcomes the permeability limitations of the commercially available molecular rotor dye, Proteostat, enabling widespread labeling of protein aggregates in live and post-fixed worms. Here, we present our novel approach and demonstrate quantification of protein aggregates in multiple tissues to detect differences in protein aggregation due to both genotype and treatment effects. We also outline anticipated future applications for this protocol, including its adaptation for other dyes, as well as limitations and special considerations for its use.

## Introduction

Protein aggregates are a diverse array of intra- and extracellular structures that contain misfolded proteins and have limited macromolecular movement [1–5]. Protein aggregation is induced by aging and cellular stress and is a hallmark of neurodegenerative diseases including Huntington’s, Alzheimer’s, and Parkinson’s [5–7]. Yet protein aggregates can be toxic or adaptive, and each year new research sheds light on the causes and consequences of protein aggregation as well as the processes that mitigate toxicity [1, 8]. Generally speaking, organisms are thought to be healthier when aggregation is minimized. Therefore, tools to visualize protein aggregates in whole organisms are important for screening genes and compounds that may reduce or alter protein aggregation in connection with improved health outcomes.

Visual detection of protein aggregates *in vivo* has primarily been achieved by fluorescent protein tags or antibodies, including polyglutamine (polyQ) proteins, luciferase, and the recently developed AgHalo and AggTag systems, among others [9–17]. These methods require targeting known components of protein aggregates and are therefore limited to the type(s) of protein aggregates in which that target is found. Furthermore, fluorescent tags can alter the templating and binding of amyloid-forming species due to their disproportionate size [18–21]. The use of fluorescent probes to detect protein aggregates in live samples overcomes these limitations but is less common, particularly in whole organisms [2, 22–24]. A key barrier has been the low permeability of cells and tissues to such probes. Nevertheless, it is widely recognized that probes for detecting protein aggregates *in vivo* are an essential part of the toolkit for studying protein homeostasis [2, 24–28].

The commercially available probe, PROTEOSTAT® (referred to herein as Proteostat), is a molecular rotor dye whose fluorescence intensity increases upon binding to protein aggregates [29]. Mechanistically, the rotor component spins freely in solution so that the dye exhibits non-radiative decay. Interactions with low-fluidity structures stabilize the rotor, permitting fluorescence. This rational design was based on the mechanism of a commonly used but dimmer dye, ThT [29]. Since its introduction, Proteostat has been used extensively as a probe for flow cytometry and visual detection of protein aggregates in mammals, insects, nematodes, bacteria, and plants [30–39]. Its peak excitation is ∼470 nm, and it emits between 550 and 690 nm, making it accessible for standard confocal imaging systems with a 488 nm laser [40]. Despite all of this, its use has been extremely limited in whole organism samples, perhaps due to inaccuracies in the literature [34].

Over the past decade, there has been a widely held misconception that Proteostat requires fixation and so cannot be used to stain live tissue [11, 17, 23, 29, 41–43]. This cannot be the case, as living cells have been labeled effectively with Proteostat for flow cytometry [37, 44, 45]. Additionally, it was used to visualize aggregates in live cell culture with active *Plasmodium* infection [32]. The misconception likely arose from the manufacturer’s guidance that the Proteostat Aggresome Detection Kit is not optimized for use in live cells [40]. However, failures to stain live cells and tissues are more likely due to low permeability in live as opposed to fixed, permeabilized tissues. In the meantime, it has been recognized that non-specific labeling of protein aggregates in living tissue is critically important for tracking the dynamics of protein aggregates under physiological conditions. Therefore, several other dyes have recently been developed to label aggregates in living tissue, though none are yet as readily available as Proteostat [17, 23, 26, 41–43, 46, 47]. Additionally, while many of them have enhanced membrane permeability, they have yet to be applied in whole organisms.

*C. elegans* is a popular and convenient model organism for the study of protein aggregation mechanisms in an intact animal during normal development, disease, and aging [14, 34, 48–60]. However, permeability barriers have largely limited the study of protein aggregation to endogenously labeled, aggregation-prone proteins and peptides. To achieve visualization of non-specific protein aggregates in whole *C. elegans* nematodes without the need for strain engineering, we have developed a simple, inexpensive soaking protocol that overcomes the permeability limitations of the Proteostat dye, enabling visualization in post-fixed or living tissue throughout the worm body. Here, we demonstrate the use of this protocol to label and quantify protein aggregates in multiple tissues, detecting differential protein aggregation due to genotype and treatment effects. We also outline anticipated future applications for this protocol, including its adaptation for other dyes, as well as limitations and special considerations for its use.

## Methods

### Full Protocol

A full protocol is published at https://dx.doi.org/10.17504/protocols.io.261gek1k7g47/v1.

### C. elegans Strains and Maintenance

*C. elegans* strains were maintained on nematode growth (NGM) plates seeded with OP50 *E. coli* at 20°C [61].The wild type reference strain used in this study is N2 obtained from the C*aenorhabditis* Genetics Center in 2019 and bearing *alh-2(ot588)*. AM44 (*rmIs172 [F25B3.3p::Q67::CFP]*), AM49 (*rmIs172 [F25B3.3p::Q19::CFP]*), VC1099 (*hsp-4(gk514)* II.), VC475 (*hsp116.2(gk249)* V.), and RB1098 (*hsp-12.6(ok1077)* IV.) were obtained from the *Caenorhabditis* Genetics Center. AM44 and AM49 were a gift from the Morimoto Lab. RB1098, VC1099, and VC147 were created by the *C. elegans* Gene Knockout Consortium [62].

### Proteostat Staining

Proteostat staining was conducted before proteotoxic stress treatment for live imaging or after proteotoxic stress treatment and recovery for fixed imaging. Briefly, worms were washed in M9 buffer, resuspended in M9 to a volume of 100 μL and exposed to 0.6% v/v PROTEOSTAT® Aggresome Detection Reagent (Enzo, Cat. #ENZ-PADR-A10C-0000) with 3 × 10^8^ cfu/mL of live *E. coli* OP50 culture in M9 buffer or vehicle (3 × 10^8^ cfu/mL of live *E. coli* OP50 culture in M9) for 2–3 hours on a benchtop shaker at 20-21°C. Worms were washed 3 times with M9, transferred to seeded NGM plates, placed in a drawer to block out light, and allowed to “rest” for 1–2 hours at 20-21°C. After the rest period, worms were washed twice with M9 and immediately fixed or prepared for live imaging.

### Acute Heat Stress

To induce rapid protein aggregation for live imaging, populations of synchronized worms were washed four times for 15 min to clear gut bacteria, stained with Proteostat, washed three times for 10 min, and then exposed to 35°C (heat stress) or 20°C (controls) for 20 min while suspended in M9 buffer in a heat block. Worms were then allowed to rest and recover for 2 hours on seeded NGM plates at room temperature before washing of excess bacteria, mounting on agarose pads, and live imaging.

### Tunicamycin Stress

Synchronized populations of young adults (200–400 worms per plate per strain) were washed into tubes with M9. Worms were exposed to 7.33 μg/mL tunicamycin (Cell Signaling Technology Cat. #12819) with 3 × 10^8^ cfu/mL of live *E. coli* OP50 culture in M9 or to vehicle (3 × 10^8^ cfu/mL of live *E. coli* culture in M9) for 3 hours on a benchtop shaker at room temperature (20-21°C). After the exposure, worms were washed 3 times with M9, plated onto NGM plates seeded with *E. coli* OP50, and allowed to recover for 17 hours before Proteostat staining, rest, and fixation.

### Hypoxia Stress

Synchronized populations of young adults on NGM plates seeded with *E. coli* OP50 were exposed to “control” or “hypoxia” conditions on seeded NGM plates followed by recovery, Proteostat staining, rest, and fixation. Prior to stress exposure, 100 mM CuSO_4_ was spread around the perimeter of the agar on each plate to discourage worms from crawling up the sides of the plates. Hypoxic conditions were achieved in a hypoxia chamber at room temperature connected to a compressed gas cylinder filled with 99.9% N_2_ and 0.1% O_2_. A fiber-optic Firesting O_2_ sensor and temperature probe from PyroScience were placed in the chamber to continuously monitor pO_2_ and temperature inside the chamber throughout the experiment. Oxygen and temperature data were recorded using Pyro Workbench software. Oxygen remained stable between 0.1–0.2% for all experiments, and temperature was stable between 20 and 21°C. “Hypoxia” samples were kept under hypoxic conditions for 24 hours followed by 18 hours of recovery in room air at 20-21°C. Because of nearly complete developmental arrest during hypoxia exposure, “control” samples were exposed to 0 hours of hypoxia followed by 18 hours of mock recovery at 20-21°C.

### Fixation and Mounting

For fixed imaging, worms were fixed in 4% PFA (Paraformaldehyde, 32% w/v aq. Soln, ThermoFisher Cat. #47377) with 0.2% v/v Triton X-100 (ThermoFisher Cat. #AC215680000) in M9 buffer for 30 minutes at 4°C and then washed three times in fresh M9. Excess buffer was removed from each tube, leaving ∼30 μL of volume behind, and then 10 μL of 100% glycerol was added as a mounting medium. The worms were applied to a glass slide, a cover slip was placed on top, and the slides’ edges were sealed with nail polish. Slides were stored at 4°C for up to 1 week before imaging. For live imaging, a few drops of 4 mM levamisole (Levamisole Hydrochloride, Sigma-Aldrich, Cat. #1359302-125MG) were added to 4% agarose pads on microscope slides. Worms were immediately transferred to the levamisole drop, petroleum jelly was spread around the outside of the agar pad, and a cover slip was placed on top. Worms were imaged live within 1 hour of mounting.

### Imaging

Z-stacks through Proteostat-stained live or fixed worms were collected on a Leica TCS SP8 Laser Scanning Confocal Microscope with an OPSL 488 laser set to 9% or 10% power and a HC PL APO CS2 40×/1.30 oil objective. The detector collection window was set between 500 and 710 nm for all experiments except colocalization experiments. For colocalization experiments, the collection window was changed to 595–700 nm to avoid overlap with CFP collection (500–575 nm).

### Quantification of Protein Aggregates using ImageJ

Proteostat puncta were quantified using ImageJ software. Z-stacks of individual worms were collapsed into maximum intensity projections. A region of interest was cropped out of the projection, and a threshold was set to subjectively maximize the detection of Proteostat puncta while minimizing the detection of background. Threshold settings were determined from the images in an experiment with the highest dynamic range and kept constant for all other images in the experiment. In the Set Measurements menu, Area, Min & Max Gray Value, Integrated Density, Standard Deviation, and Mean Gray Value were checked. Areas above threshold within the ROI were then measured using the Analyze Particles tool. The summary data, which included the number of particles, the total area occupied by particles, the average size of the particles, and the integrated density of the particles in the ROI, was obtained from the software and recorded. To quantify punctate vs. diffuse patterns in Fig. 5, average fluorescence intensity and standard deviation were measured across a standardized 60×60 pixel ROI within maximum intensity projections and used to calculate the coefficient of variation 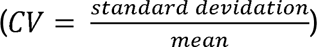.

### Statistical Analysis

Statistical analyses were performed using R Statistical Software (R.app GUI 1.82-8536 High Sierra build, RStudio 2026.01.1+403) to build generalized linear models. Data were fitted to GLMs with either gaussian or negative binomial (nbinom2) distributions using the glmmTMB package v.1.1.13 [63]. Residual diagnostics were performed using the DHARMa package v.0.4.7 according to the developer’s guidelines [64]. When model selection was warranted, the minimum Bayesian Information Criterion (BIC) was used to select the optimal model. The emmeans package v. 1.11.2-8 was used to calculate estimated marginal means, standard errors, and Tukey-corrected p-values for pairwise contrasts [65]. Other R packages used for data manipulation and plotting included tidyverse v.2.0.0, ggbeeswarm v.0.7.2, and the most up-to-date dependencies for all the above listed packages[66].

## Results

### Proteostat and polyQ::CFP signals colocalize in neural aggregates

Despite just one previous study having used Proteostat dye on fixed *C. elegans* tissues [34], we set out to develop a soaking protocol to label aggregates in live or fixed, whole *C. elegans* nematodes. A key challenge in staining with Proteostat, as with other exogenous fluorescent probes, antibodies, and drugs, is the uptake of the dye past the worm cuticle and into the deeper tissues via liquid ingestion or soaking. In fact, even axenic culture of *C. elegans* was notoriously challenging because the worms would not ingest sufficient nutrients while suspended in liquid [67, 68]. A breakthrough for axenic culture came from a study that established that *C. elegans* require suspended particles roughly the size of bacterial cells to stimulate swallowing [67]. The authors demonstrated that inclusion of even abiotic particles of the proper size in axenic media is sufficient to allow growth and development comparable to media containing actual bacterial food. With this in mind, we included a small amount of bacterial food in the staining solution to encourage ingestion of the dye. This permitted the use of a simple soaking protocol to generate a heat-shock induced increase in Proteostat signal in live worms (Fig. 1a, b). Briefly, synchronized young adults were exposed to Proteostat dye prior to a 20-minute 35°C heat shock and then allowed to “rest” and “recover” (see Methods) with food for 1 hour prior to live imaging. To confirm that the signal was present in protein aggregates, we performed the same protocol on strain AM44, which expresses polyQ67::CFP in neurons that incorporates into aggregates after heat stress [49]. We found Proteostat signal present in various structures throughout the head after heat stress, including in neuron cell bodies that were positive for bright polyQ67::CFP signal, confirming that Proteostat colocalizes with a known component of protein aggregates (Fig. 1c, Online Resource 1). Proteostat signal was also apparent in many other tissues throughout the head (Fig.1c, Online Resource 1).

**Fig. 1.**
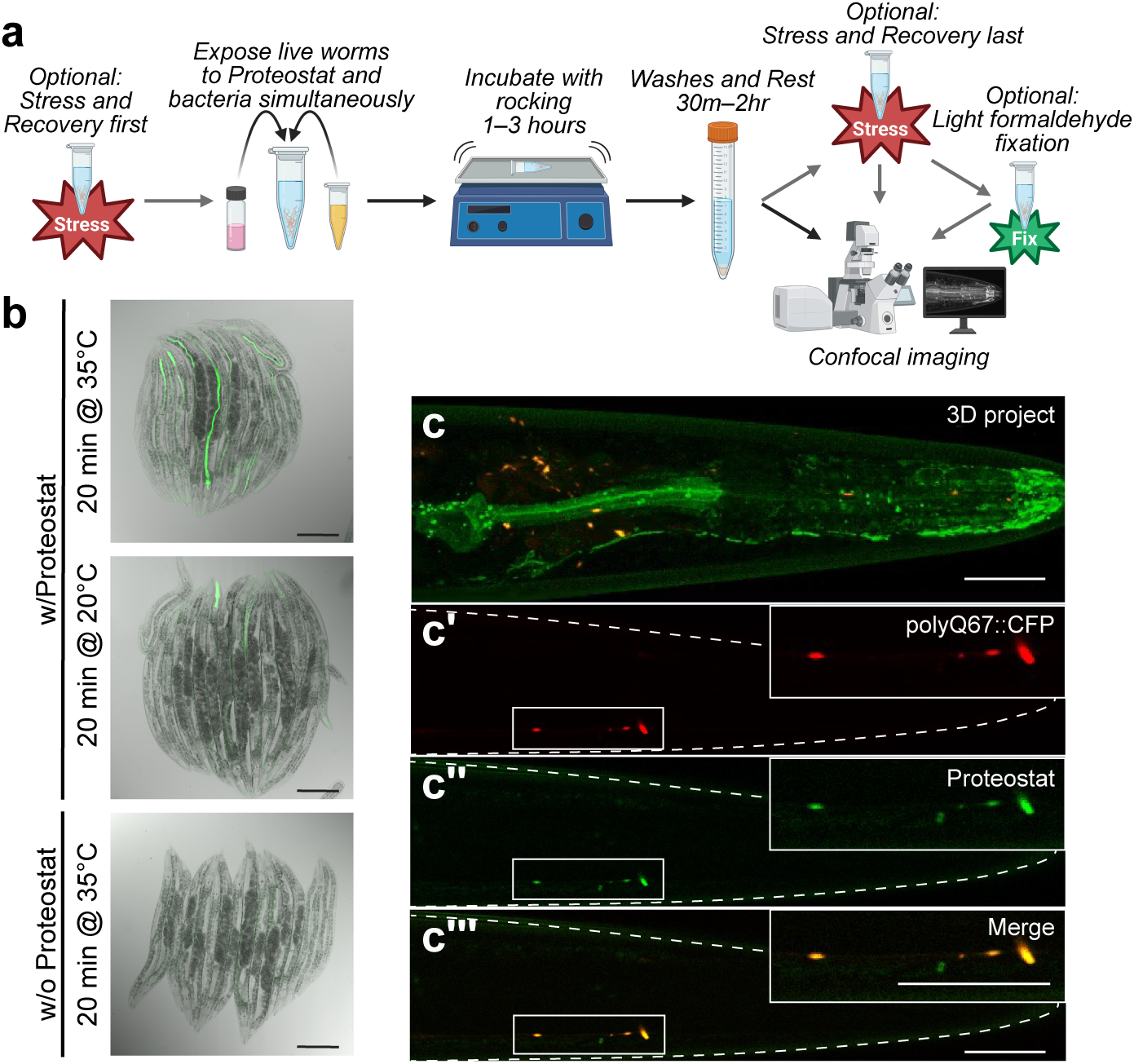
Proteostat and polyQ::CFP signals colocalize in neural aggregates. a) Schematic of the new Proteostat staining protocol for *C. elegans*. Black arrows denote mandatory steps. Gray arrows denote optional steps. b) Live imaging shows Proteostat signal (green) increases after a 20-minute heat shock at 35°C. Scale bar = 200 µm. c) 3D projection of a Z-stack through the head region showing colocalization of polyQ67::CFP in neurons (red) and Proteostat (green) in live worms after heat stress. c’–c’’’) A single slice from the Z-stack in (c). Scale bars = 20 µm. The schematic in (a) was created in BioRender. Fausett, S. (2026) https://BioRender.com/ksvdfbt

### Multiple proteotoxic stresses increase Proteostat signal in anterior tissues

Protein aggregation is known to increase after ER or hypoxia stress [52, 69]. Therefore, we tested the ability of Proteostat to report increased protein aggregation after these proteotoxic stresses. To induce ER stress, early adults were exposed in liquid to 7.33 µg/mL tunicamycin or M9 buffer (vehicle) for 3 hours followed by 17 hours of recovery, Proteostat staining, rest, fixation, and imaging. Tunicamycin treatment increased Proteostat signal in the buccal cuticle, including the flaps and the collar. These structures were previously shown to contain high levels of intrinsically disordered and prion-like proteins and stain positively for Congo Red and Thioflavin T, dyes that bind amyloid sheets (Online Resource 2, Fig. 2a, arrowheads) [70, 71]. Additionally, bright puncta were visible in other tissues of the anterior head, the isthmus, the terminal pharyngeal bulb, and the excretory cell (Online Resource 2, Fig. 2a, b). In contrast, worms treated with tunicamycin but not exposed to Proteostat showed no visible puncta (Fig. 2c). To quantify puncta in the anterior head, we captured Z-stacks through the head, cropped from the oral aperture to the posterior aspect of the anterior pharyngeal bulb (dashed outlines in Fig. 2a, b), and collapsed the stacks to maximum intensity projections. Using the Analyze Particles tool, we quantified the Total Area, Count, and Integrated Density of the puncta in ImageJ to show a tunicamycin-induced increase in Proteostat puncta within this anterior region (Fig. 2d–f). We also quantified puncta in the excretory cell body to reveal significantly increased signal in the tunicamycin-treated worms (Fig. 2g–i). We obtained similar results when we exposed early adults to 24 hours of 0.15% O_2_ hypoxia stress followed by 18 hours of recovery, Proteostat staining, rest, fixation, and imaging. We saw increased Proteostat signal using all measurement methods in the buccal cuticle, pharyngeal epithelium, head muscles, neurons, and excretory cell, among other tissues (Fig. 3). Together, these data strongly suggest that Proteostat faithfully reports increased protein aggregation in anterior tissues after known proteotoxic stresses.

**Fig. 2.**
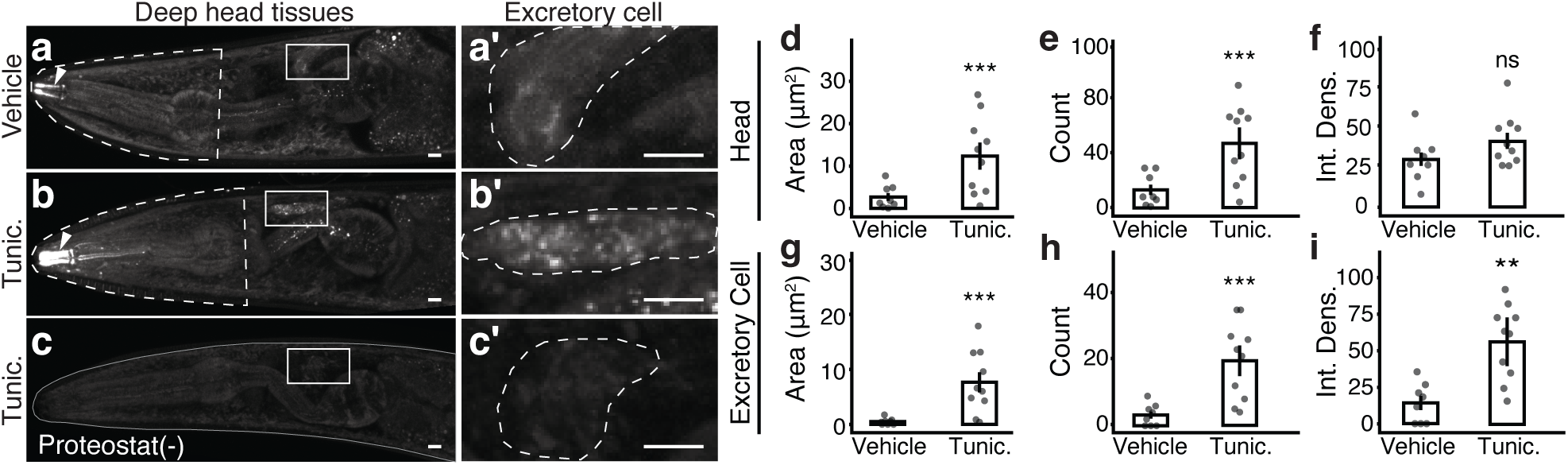
Tunicamycin increases Proteostat signal in anterior tissues. a–c) Young adults were exposed to 7.33 µg/mL tunicamycin or M9 vehicle for 3 h followed by 17 h recovery, Proteostat staining, rest, fixation, and imaging. For the head tissues, Z-stacks were collected from the top of the nerve ring to the bottom of the anterior pharyngeal bulb. The ROI was drawn from the oral aperture to the posterior aspect of the anterior pharyngeal bulb, following the outline of the head. Arrowheads indicate amyloid structures in the buccal cuticle. a’–c’) For the excretory cells, Z-stacks were collected from the top of the excretory cell body to the bottom, and the cell was outlined in a maximum intensity projection to generate the ROI (white dashed outline). c, c’) A Proteostat negative control worm outlined in solid white. Z-stacks were collected from the top of the nerve ring to the bottom of the first pharyngeal bulb. Scale bars = 5 µm. d–i) The Analyze Particles tool in ImageJ was used to quantify the Total Area, Count, and Integrated Density of particles above a common threshold in maximum intensity projections of the ROI. ***p<0.001, **0.001<p<0.01, *0.01<p<0.05. Statistical data are provided in Online Resource 3–8

**Fig. 3.**
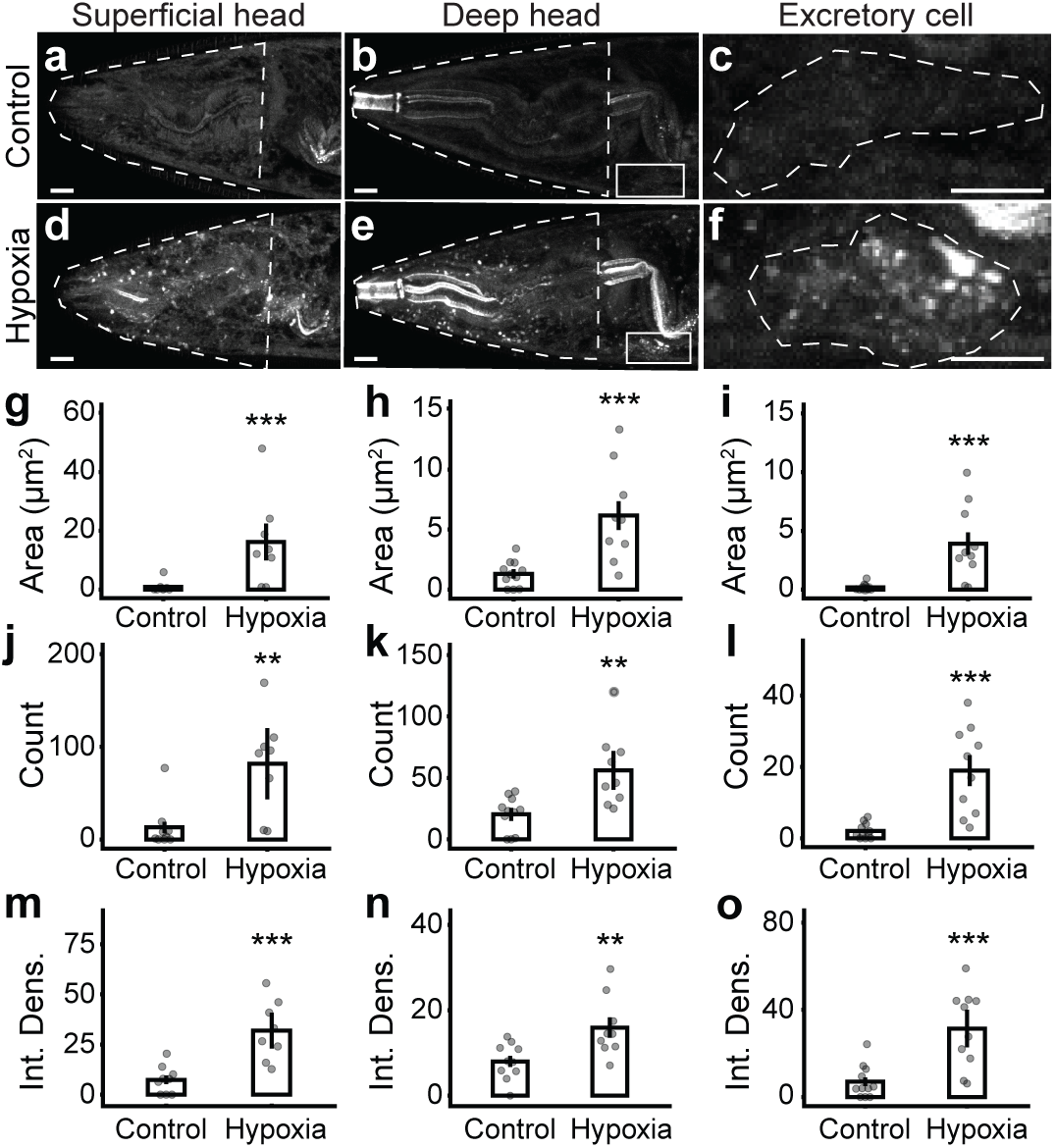
Hypoxia increases Proteostat signal in anterior tissues. a–f) Young adults were exposed to 24 h 0.15% O_2_ hypoxic or 0 h hypoxic (control) conditions followed by 18 h recovery. After recovery, samples were processed for Proteostat staining, rest, fixation, and imaging. ROI for quantification are marked by white dashed outlines. For “superficial head” tissues, Z-stacks were collected from the top of the worm to the top of the anterior pharyngeal bulb. For “deep head” tissues, Z-stacks were collected from the top of the nerve ring to the bottom of the anterior pharyngeal bulb. The ROI for both (white dashed outlines) was drawn from the oral aperture to the posterior aspect of the anterior pharyngeal bulb, following the outline of the head. For the excretory cells, Z-stacks were collected from the top of the excretory cell body to the bottom, and the cell was outlined to generate the ROI (white dashed outlines). Scale bars = 5 µm. g–o) The Analyze Particles tool in ImageJ was used to quantify the Total Area, Count, and Integrated Density of particles above a common threshold in maximum intensity projections of the ROI. Graphs correspond to the images directly above them. ***p<0.001, **0.001<p<0.01, *0.01<p<0.05. Statistical data are provided in Online Resource 9–17

### Tunicamycin-induced Proteostat signal in the excretory cell is dependent on the HSP70 ortholog, HSP-4

We next decided to test whether Proteostat signal responds to genetic manipulations known to impact protein homeostasis. Previously, it was shown that worms lacking the HSP70 ortholog, *hsp-4*, have enhanced autophagy and tolerance to proteotoxic stress [72]. We, therefore, assessed whether Proteostat signal is reduced in the excretory cells of tunicamycin-treated *hsp-4(gk514)* adults. Indeed, we found that while the integrated density of Proteostat signal was increased by tunicamycin treatment in the N2 reference strain, it was significantly reduced in the *hsp-4(gk514)* mutants (Fig. 4), supporting the hypothesis that Proteostat puncta may be cleared by elevated autophagy in these mutants.

**Fig. 4.**
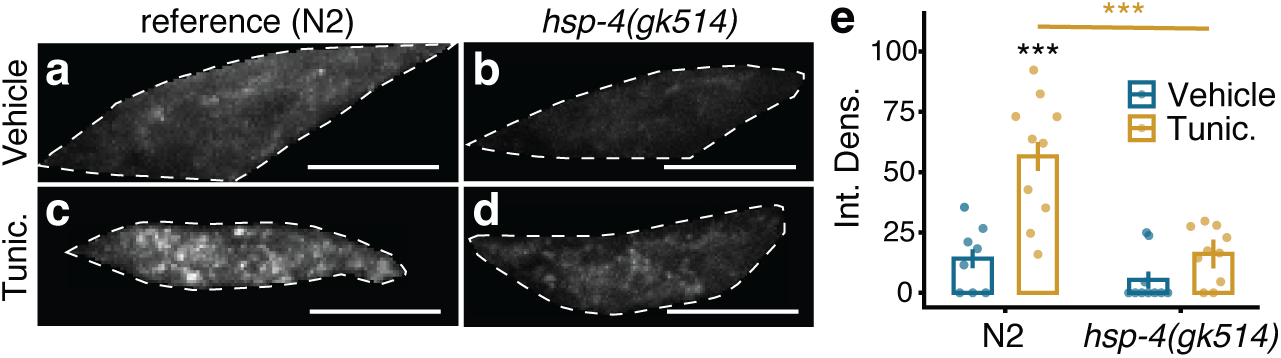
Tunicamycin-induced Proteostat signal in the excretory cell is dependent on HSP-4. a–d) Young adults were exposed to 7.33 µg/mL tunicamycin or M9 (vehicle) for 3 h followed by 17 h recovery, Proteostat staining, fixation, and imaging. Z-stacks were collected from the top of the excretory cell body to the bottom, and the cell was outlined to generate the ROI (white dashed outline). Scale bars = 5 µm. e) The Analyze Particles tool in ImageJ was used to quantify the Integrated Density of particles above a common threshold in maximum intensity projections of the ROI. ***p<0.001, **0.001<p<0.01, *0.01<p<0.05. Statistical data are provided in Online Resource 18

### Proteostat reveals differential effects of sHSPs on hypoxia-induced protein aggregation among posterior tissues

Small heat shock proteins (sHSPs) bind and sequester misfolded proteins in phase-separated compartments to stabilize the proteome under proteotoxic stress [4, 5]. These structures appear similar to more solid inclusions under low resolution and may transition to solid aggregates. In theory, Proteostat could label both categories of aggregates. To begin exploring the effects of sHSPs on the appearance of the Proteostat foci, we exposed *hsp-12.6(ok1077)* and *hsp-16.2(gk249)* mutants to hypoxia stress. Interestingly, we observed differential effects among posterior tissues. Hypoxia stress increased the integrated density of Proteostat signal in the tissues surrounding the rectum in the N2 reference strain and in *hsp-16.2(gk249)* mutants (Fig. 5a–g). Meanwhile, the absence of *hsp-12.6*, but not *hsp-16.2*, increased the Proteostat signal in rectal tissues compared to N2 in both the hypoxia-treated and control groups (Fig. 5a–h). Surprisingly, in the nearby posterior intestine, puncta that were readily apparent in all genotypes under control conditions were abolished and replaced by a more diffuse signal in sHSP mutants, but not the reference strain, after hypoxia stress (Fig. 5a–f). To quantify this effect, we isolated a rectangular region of interest from maximum intensity projections inside the posterior intestine just anterior to the rectum (Fig. 5a–f, pink ROI) and measured both the mean fluorescence intensity and the coefficient of variation. While the mean intensity of fluorescence was similar across genotypes and treatments (Fig. 5i), the coefficient of variation was reduced in hypoxia-treated worms, indicating a significantly less punctate pattern (Fig. 5j). This effect was more pronounced in the sHSP mutants than in the N2 reference strain. Altogether, these observations suggest that sHSPs may play different roles in distinct tissues to organize misfolded proteins under stress. Our results further demonstrate that Proteostat is an excellent tool for observing differential protein aggregation patterns simultaneously across multiple tissue types.

**Fig. 5.**
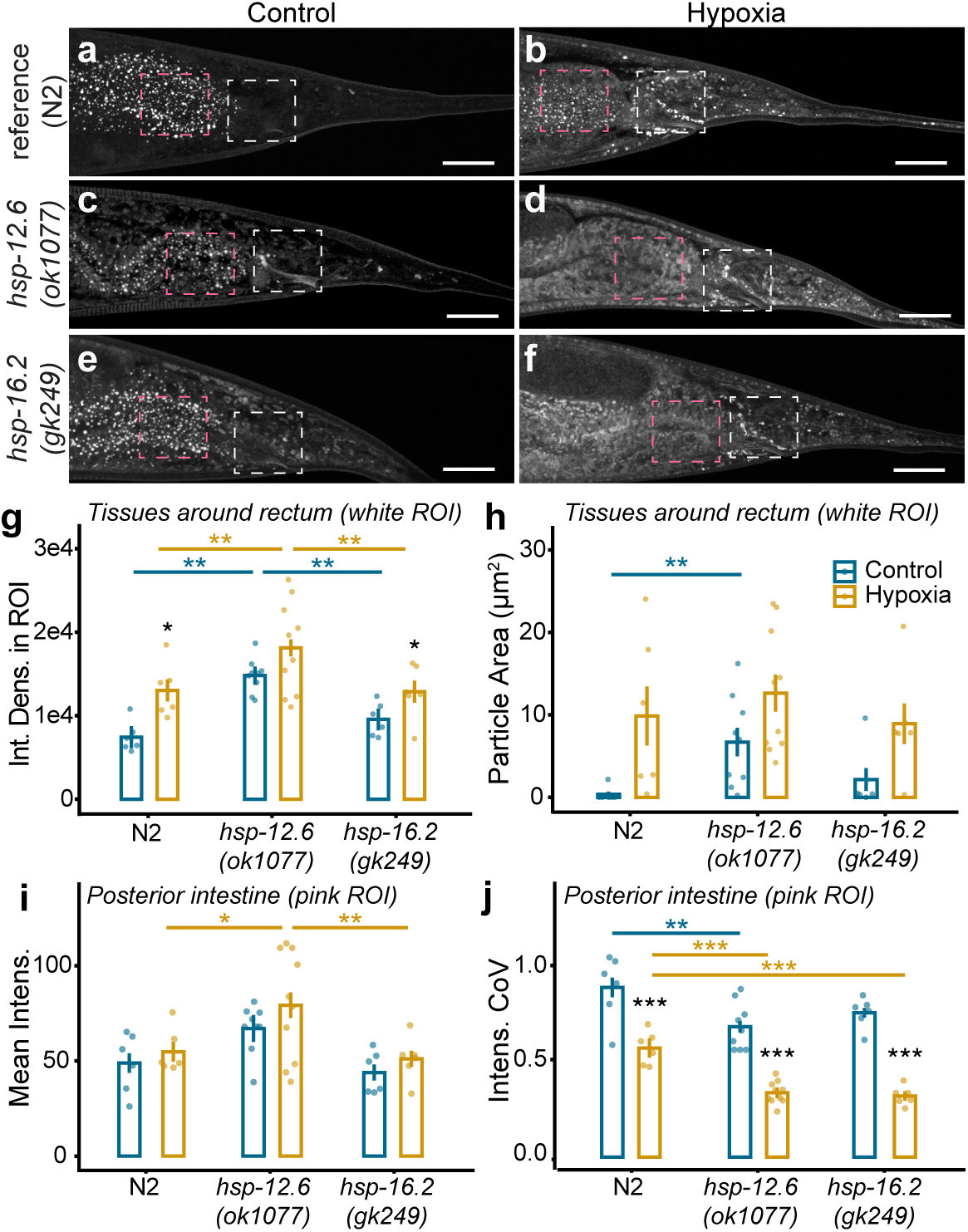
Proteostat reveals differential effects of sHSPs on hypoxia-induced protein aggregation among posterior tissues. a–f) Young adults were exposed to 24 h 0.15% O_2_ hypoxic or 0 h hypoxic (control) conditions followed by 18 h recovery. After recovery, samples were processed for Proteostat staining, rest, fixation, and imaging. Two ROI were selected for imaging, one including the tissue surrounding the rectum and anus (white dashed outlines) and one in the most posterior intestinal cell (pink dashed outlines). Scale bars = 20 µm. g, h) For the rectal tissue, Integrated Density of fluorescence was calculated for the entire ROI (g), and Analyze Particles in ImageJ was used to determine the Total Area occupied by particles above a common threshold within the ROI (h). i, j) For the intestine, mean pixel intensity (I) and standard deviation were measured for the pink ROI and used to calculate the coefficient of variation (J) to quantify the evenness of the signal. ***p<0.001, **0.001<p<0.01, *0.01<p<0.05. Statistical data are provided in Online Resource 19–22

## Discussion

### An accessible, dye-based protocol for detecting protein aggregates in living or fixed nematodes

Historically, the use of fluorophore-tagged polyQ peptides to detect protein aggregation has been very successful in the worm [3, 14, 16, 49, 50, 55, 73–77]. However, these tools detect only a single component of protein aggregates, potentially missing certain aggregate types. They also fluoresce diffusely when not incorporated into aggregates, resulting in high background signal that may obscure tiny, more diffuse aggregates. Finally, these tools require genetic engineering or genetic crosses to quantify protein aggregation across genetic backgrounds and are usually expressed in a specific tissue. Proteostat dye allows the detection of any structure that is viscous enough to prevent resonance transfer in the rotor dye, and it does not fluoresce unless incorporated into these structures, resulting in exceptionally low background. Furthermore, it reports protein aggregation throughout the worm and does not require any genetic manipulation, providing an entry point for high-throughput applications such as screening. In our study, most tissues except for the germline were seen to take up the dye, especially in the presence of proteotoxic stress (Fig. 2–5). Although we rarely saw Proteostat signal in the germline, we suspect this reflects a feature of active germline tissue rather than a lack of dye penetrance. Future work should establish whether conditions known to produce protein aggregates in the germline, such as oocyte arrest, also promote Proteostat puncta with this protocol [34].

### Recovery and rest periods are important for detecting aggregates with Proteostat

During troubleshooting, we discovered that recovery periods and rest periods were critical components of our successful attempts. “Recovery periods” refer to recovery time after proteotoxic stress (presumably allowing aggregates to develop), and “rest periods” refer to a short period during which Proteostat dye is allowed to incorporate into aggregates and clear from the gut lumen. In general, the length of the recovery period should be optimized for a given stress and to suit the needs of the experiment. For instance, in our hypoxia protocol, worms were exposed to hypoxia for 24 hours then allowed to recover overnight (∼18 hours) before Proteostat staining. After staining, the worms were allowed a rest period of 1–2 hours on plates before imaging or fixation. Including these steps dramatically increased Proteostat signal but meant that aggregation could not be assessed during or immediately after stress exposure. To overcome this limitation for heat stress, we exposed the worms to Proteostat prior to heat stress so that the rest period overlapped with the stress exposure. This approach was successful and demonstrates the flexibility of our protocol.

### Proteostat dye is effective in live tissue

Several sources claim that Proteostat (specifically the Aggresome Detection Reagent) can only be used in fixed cells and tissues [11, 17, 23, 29, 41–43]. We believe this misconception has unfortunately prevented its optimization and more widespread use. Only one previous study has used Proteostat in *C. elegans*, employing an effective but cumbersome protocol that involves prestaining bacterial food and/or loading the agar plates with large amounts of dye before allowing worms to ingest that food over hours to days [34, 78]. Additionally, the authors fixed their samples prior to imaging, so it remained unclear whether Proteostat dye could report aggregates in living tissue. Regardless, we suspected that the past failures to stain live tissues were due to dye penetrance, rather than some other fixation effect, since fixation usually involves a permeabilization step. Our work clearly shows that Proteostat can be used to image aggregates in live tissue. Our new method requires a relatively low concentration of dye, can be executed over just a few hours, and overcomes the permeability limitation by including bacterial food to stimulate the worms’ swallowing behavior in liquid. Since abiotic bacteria-sized particles also stimulate swallowing [67], our protocol could be adapted for starvation stress by including abiotic particles of the appropriate size in place of bacteria. However, since ingestion of the dye itself is presumably required, our protocol would not be expected to stain dauer larvae or embryos, which do not feed. An exciting implication of our work is that other poorly penetrating dyes could be used more effectively if bacteria are simply included in the staining solution, potentially enabling a variety of dye-based kinetic studies in the worm. Although Proteostat could in theory report real-time protein aggregation kinetics, its nature as a molecular rotor dye still makes this challenging: it has only moderate photostability and requires low-wavelength excitation. While these properties might limit its usefulness for long-term kinetic studies of protein aggregation, short-term time-lapse is likely possible under optimized imaging conditions.

## Supporting information

Online Resource 1

Online Resource 2

## Acknowledgements

We are grateful to Richard Morimoto and colleagues for providing the AM44 and AM49 lines to the community and the *C. elegans* Gene Knockout Consortium for generating alleles used in this study. Strains were maintained and distributed by the CGC, which is funded by NIH Office of Research Infrastructure Programs (P40 OD010440). For funding, we thank the University of North Carolina Wilmington, generous donations from the Cavalieri family to support undergraduate research, and the UNCW Center for the Support of Undergraduate Research & Fellowships. Finally, we thank the Richard M. Dillaman Bioimaging Facility for use of instruments and technical support.

## Author Contributions

Conceptualization: Sarah Fausett and Tarah Krishingenr; Optimization and Experimentation: Tarah Krishingner; Data analysis and figure development Sarah Fausett; Writing - original draft preparation: Tarah Krishingner and Sarah Fausett; Writing - review and editing: Tarah Krishingner and Sarah Fausett; Funding acquisition: Sarah Fausett; Resources: Sarah Fausett; Supervision: Sarah Fausett.

