## Supplemental Information for "A New Soaking Protocol to Live Stain *C. elegans* with Proteostat Dye Reveals Global Patterns of Stress-Induced Protein Aggregation"

Tarah Krishingner ([0009-0009-2819-2327](https://nam12.safelinks.protection.outlook.com/?url=https%3A%2F%2Forcid.org%2F0009-0009-2819-2327&data=05%7C02%7Cfausetts%40uncw.edu%7Ca63f3a0e58014e29e04108df0f882b63%7C2213678197534c75af2868a078871ebf%7C0%7C0%7C639246750009544552%7CUnknown%7CTWFpbGZsb3d8eyJFbXB0eU1hcGkiOnRydWUsIlYiOiIwLjAuMDAwMCIsIlAiOiJXaW4zMiIsIkFOIjoiTWFpbCIsIldUIjoyfQ%3D%3D%7C0%7C%7C%7C&sdata=KiMTordmMYw5JMOrSNBQvfVlfxdInDGdfByPn6kO1pI%3D&reserved=0))^1#a^ & Sarah R. Fausett ([0000-0003-2046-0226](https://orcid.org/0000-0003-2046-0226))^1#a^

**This PDF file includes:**

Legend for Source Data (separate Excel file)

Legend for Online Resource 1 (.mp4 video file)

Legend for Online Resource 2 (.mp4 video file)

Online Resources 3 – 22 (statistical tables)

Legend for Source Data (separate .xlsx file)

**Each of the items listed is a separate sheet in Source Data.xlsx**

**Fig. 2d–f:** Young adults were exposed to 7.33 µg/mL tunicamycin or M9 vehicle for 3 h followed by 17 h recovery, Proteostat staining, rest, fixation, and imaging. Z-stacks were collected from the top of the nerve ring to the bottom of the anterior pharyngeal bulb. The ROI was drawn from the oral aperture to the posterior aspect of the anterior pharyngeal bulb, following the outline of the head. The Analyze Particles tool in ImageJ was used to quantify the Area, Count, and Integrated Density of particles above a common threshold in maximum intensity projections of the ROI. Accompanies Fig. 2d–f. Statistical data are provided in Online Resource 3–5

**Fig. 2g–i:** Young adults were exposed to 7.33 µg/mL tunicamycin or M9 vehicle for 3 h followed by 17 h recovery, Proteostat staining, rest, fixation, and imaging. Z-stacks were collected from the top of the excretory cell body to the bottom, and the cell was outlined to generate the ROI. The Analyze Particles tool in ImageJ was used to quantify the Area, Count, and Integrated Density of particles above a common threshold in maximum intensity projections of the ROI. Accompanies Fig. 2g–i. Statistical data are provided in Online Resource 6–8

**Fig. 3g, j, m:** Young adults were exposed to 24 h 0.15% O_2_ hypoxic or 0 h hypoxic (control) conditions followed by 18 h recovery, Proteostat staining, rest, fixation, and imaging. For “superficial head” tissues, Z-stacks were collected from the top of the worm to the top of the anterior pharyngeal bulb. The ROI was drawn from the oral aperture to the posterior aspect of the anterior pharyngeal bulb, following the outline of the head. Analyze Particles in ImageJ was used to determine the Total Area, Count, and Integrated Density of particles above a common threshold. Statistical data are provided in Online Resource 9–11

**Fig. 3h, k, n:** Young adults were exposed to 24 h 0.15% O_2_ hypoxic or 0 h hypoxic (control) conditions followed by 18 h recovery, Proteostat staining, rest, fixation, and imaging. For “deep head” tissues, Z-stacks were collected from the top of the nerve ring to the bottom of the anterior pharyngeal bulb. The ROI was drawn from the oral aperture to the posterior aspect of the anterior pharyngeal bulb, following the outline of the head. Analyze Particles in ImageJ was used to determine the Total Area, Count, and Integrated Density of particles above a common threshold. Statistical data are provided in Online Resource 12–14

**Fig. 3i, l, o**: Young adults were exposed to 24 h 0.15% O_2_ hypoxic or 0 h hypoxic (control) conditions followed by 18 h recovery, Proteostat staining, rest, fixation, and imaging. Z-stacks were collected from the top of the excretory cell body to the bottom, and the cell was outlined to generate the ROI. Analyze Particles in ImageJ was used to determine the Total Area, Count, and Integrated Density of particles above a common threshold. Statistical data are provided in Online Resource 15–17

**Fig. 4e:** Young adults were exposed to 7.3 µg/mL tunicamycin or M9 (vehicle) for 3 h followed by 17 h recovery, Proteostat staining, rest, fixation, and imaging. Z-stacks were taken from the top of the excretory cell body to the bottom. The excretory cell was outlined, the background was cleared from maximum intensity projections, and Analyze Particles in ImageJ was used to determine the Integrated Density of particles above a common threshold. ***p<0.001, **0.001<p<0.01, *0.01<p<0.05. Statistical data are provided in Online Resource 18

**Fig. 5g-h:** Young adults were exposed to 24 h 0.15% O_2_ hypoxic or 0 h hypoxic (control) conditions followed by 18 h recovery, Proteostat staining, rest, fixation, and imaging. For the rectal tissue, Integrated Density of fluorescence was calculated within a rectangular 60x60 pixel ROI surrounding the rectum. Analyze Particles in ImageJ was used to determine the Total Area occupied by particles above a common threshold within the ROI. Statistical data are provided in Online Resource 18–20

**Fig. 5i-j:** Young adults were exposed to 24 h 0.15% O_2_ hypoxic or 0 h hypoxic (control) conditions followed by 18 h recovery, Proteostat staining, rest, fixation, and imaging. For the intestine, mean pixel intensity (I) and standard deviation were measured for a rectangular 60x60 pixel ROI in the most posterior intestinal cell and used to calculate the coefficient of variation to assess the evenness of the signal. ***p<0.001, **0.001<p<0.01, *0.01<p<0.05. Statistical data are provided in Online Resource 21–22

**Online Resource 1 (Accompanies Fig. 1)** A video of a rotating 3D projection of Proteostat signal (green) and neuronal polyQ67::CFP signal (red) in the head of a living worm after heat shock. To induce rapid protein aggregation for live imaging, populations of synchronized young adults were washed four times for 15 min to clear gut bacteria, stained with Proteostat, and washed again. Worms were then exposed to 35°C (heat shock) or 22°C (controls) for 20 minutes while suspended in M9 buffer and then allowed to rest and recover on seeded NGM plates for 1 hour at room temperature prior to live imaging on agar pads with levamisole

**Online Resource 2 (Accompanies Fig. 2)** A video of a rotating 3D projection of Proteostat signal in the head of a fixed worm treated with tunicamycin. Young adults were exposed to 7.33 µg/mL tunicamycin or M9 vehicle for 3 h followed by 17 h recovery, Proteostat staining, rest, fixation, and imaging. Z-stacks were collected from the top of the anterior pharyngeal bulb to the bottom. The lookup table highlights the brightest pixels in black and lower intensity Proteostat signal in yellow and orange

**Online Resource 3 (Accompanies Fig. 2d)** Statistical analysis for differences in the total area occupied by Proteostat puncta in the anterior head due to tunicamycin treatment. a) Raw data were fitted to a generalized linear model. The R package, ‘emmeans’ was used to obtain estimated marginal means on the response scale (b) and pairwise contrasts (c) with Tukey corrected p-values. Raw data are provided as a Source Data file

**a) Anterior Head Total Area—Model Statistics**

| **Family:** nbinom2 (log) | | | | | | |
| --- | --- | --- | --- | --- | --- | --- |
| **Formula:** TotalArea ~ Treatment | | | | | | |
| **AIC:** 110.1 | **BIC:** 112.8 | **logLik:** -52.1 | | **-2*log(L):** 104.1 | **df. resid:** 15 | |
| **Dispersion parameter for nbinom2 family ():** 1.73 | | | | | | |
| **Conditional Model:** | | ***Estimate*** | ***SE*** | ***z-value*** | ***Pr(>\|z\|)*** | |
| **(Intercept)** | | 0.9974 | 0.3439 | 2.900 | 0.00373 | ** |
| **Treatment** tunicamycin | | 1.5162 | 0.4291 | 3.534 | 0.00041 | *** |
| **Signif. codes:** 0 ‘***’ 0.001 ‘**’ 0.01 ‘*’ 0.05 ‘.’ 0.1 ‘ ’ 1 | | | | | | |

**b) Anterior Head Total Area—Estimated Model Means**

| ***Treatment*** | ***response*** | ***SE*** | ***df*** | ***asymp.LCL*** | ***asymp.UCL*** |
| --- | --- | --- | --- | --- | --- |
| vehicle | 2.71112535 | 0.93236093 | Inf | 1.38171935 | 5.31960462 |
| tunicamycin | 12.3489011 | 3.16834371 | Inf | 7.46855340 | 20.4183262 |

**c) Anterior Head Total Area —Contrasts**

| ***contrast*** | ***ratio*** | ***SE*** | ***df*** | ***null*** | ***z.ratio*** | ***p.value*** |  |
| --- | --- | --- | --- | --- | --- | --- | --- |
| vehicle / tunicamycin | 0.219543 | 0.0941983 | Inf | 1 | -3.53374 | 0.000409716 | *** |
| **Signif. codes:** 0 ‘***’ 0.001 ‘**’ 0.01 ‘*’ 0.05 ‘.’ 0.1 ‘ ’ 1 | | | | | | | |

**Online Resource 4 (Accompanies Fig. 2e)** Statistical analysis for differences in the number of Proteostat puncta in the anterior head due to tunicamycin treatment. a) Raw data were fitted to a generalized linear model. The R package, ‘emmeans’ was used to obtain estimated marginal means on the response scale (b) and pairwise contrasts (c) with Tukey corrected p-values. Raw data are provided as a Source Data file

**a) Anterior Head Count—Model Statistics**

| **Family:** nbinom2 (log) | | | | | | |
| --- | --- | --- | --- | --- | --- | --- |
| **Formula:** Count ~ Treatment | | | | | | |
| **AIC:** 158.8 | **BIC:** 161.5 | **logLik:** -76.4 | | **-2*log(L):** 152.8 | **df. resid:** 15 | |
| **Dispersion parameter for nbinom2 family ():** 1.69 | | | | | | |
| **Conditional Model:** | | ***Estimate*** | ***SE*** | ***z-value*** | ***Pr(>\|z\|)*** | |
| **(Intercept)** | | 2.5745 | 0.2890 | 8.908 | < 2E-16 | *** |
| **Treatment** tunicamycin | | 1.2692 | 0.3806 | 3.335 | 0.000854 | *** |
| **Signif. codes:** 0 ‘***’ 0.001 ‘**’ 0.01 ‘*’ 0.05 ‘.’ 0.1 ‘ ’ 1 | | | | | | |

**b) Anterior Head Count—Estimated Model Means**

| ***Treatment*** | ***response*** | ***SE*** | ***df*** | ***asymp.LCL*** | ***asymp.UCL*** |
| --- | --- | --- | --- | --- | --- |
| vehicle | 13.1250000 | 3.79325890 | Inf | 7.44892129 | 23.1262514 |
| tunicamycin | 46.7000029 | 11.5665072 | Inf | 28.7404811 | 75.8821768 |

**c) Anterior Head Count—Contrasts**

| ***contrast*** | ***ratio*** | ***SE*** | ***df*** | ***null*** | ***z.ratio*** | ***p.value*** |  |
| --- | --- | --- | --- | --- | --- | --- | --- |
| vehicle / tunicamycin | 0.28104923 | 0.1069726 | Inf | 1 | -3.3346 | 0.0008541 | *** |
| **Signif. codes:** 0 ‘***’ 0.001 ‘**’ 0.01 ‘*’ 0.05 ‘.’ 0.1 ‘ ’ 1 | | | | | | | |

**Online Resource 5 (Accompanies Fig. 2f)** Statistical analysis for differences in the integrated density of Proteostat puncta in the anterior head due to tunicamycin treatment. a) Raw data were fitted to a generalized linear model. The R package, ‘emmeans’ was used to obtain estimated marginal means on the response scale (b) and pairwise contrasts (c) with Tukey corrected p-values. Raw data are provided as a Source Data file

**a) Anterior Head Integrated Density—Model Statistics**

| **Family:** nbinom2 (log) | | | | | | |
| --- | --- | --- | --- | --- | --- | --- |
| **Formula:** IntDen ~ Treatment | | | | | | |
| **AIC:** 152.9 | **BIC:** 155.6 | **logLik:** -73.5 | | **-2*log(L):** 146.9 | **df. resid:** 15 | |
| **Dispersion parameter for nbinom2 family ():** 7.51 | | | | | | |
| **Conditional Model:** | | ***Estimate*** | ***SE*** | ***z-value*** | ***Pr(>\|z\|)*** | |
| **(Intercept)** | | 3.4420 | 0.1437 | 23.956 | <2e-16 | *** |
| **Treatment** tunicamycin | | 0.3186 | 0.1905 | 1.672 | 0.0945 | . |
| **Signif. codes:** 0 ‘***’ 0.001 ‘**’ 0.01 ‘*’ 0.05 ‘.’ 0.1 ‘ ’ 1 | | | | | | |

**b) Anterior Head Integrated Density —Estimated Model Means**

| ***Treatment*** | ***response*** | ***SE*** | ***df*** | ***asymp.LCL*** | ***asymp.UCL*** |
| --- | --- | --- | --- | --- | --- |
| vehicle | 31.2495021 | 4.48991065 | Inf | 23.5799538 | 41.4136258 |
| tunicamycin | 42.9723958 | 5.37447844 | Inf | 33.6303476 | 54.9095365 |

**c) Anterior Head Integrated Density —Contrasts**

| ***contrast*** | ***ratio*** | ***SE*** | ***df*** | ***null*** | ***z.ratio*** | ***p.value*** |  |
| --- | --- | --- | --- | --- | --- | --- | --- |
| vehicle / tunicamycin | 0.7271994 | 0.138523 | Inf | 1 | -1.6723 | 0.0944643 | . |
| **Signif. codes:** 0 ‘***’ 0.001 ‘**’ 0.01 ‘*’ 0.05 ‘.’ 0.1 ‘ ’ 1 | | | | | | | |

**Online Resource 6 (Accompanies Fig. 2g)** Statistical analysis for differences in the total area occupied by Proteostat puncta in the excretory cell due to tunicamycin treatment. a) Raw data were fitted to a generalized linear model. The R package, ‘emmeans’ was used to obtain estimated marginal means on the response scale (b) and pairwise contrasts (c) with Tukey corrected p-values. Raw data are provided as a Source Data file

**a) Excretory Cell Total Area—Model Statistics**

| **Family:** nbinom2 (log) | | | | | | |
| --- | --- | --- | --- | --- | --- | --- |
| **Formula:** TotalArea ~ Treatment | | | | | | |
| **AIC:** 81.1 | **BIC:** 83.8 | **logLik:** -37.6 | | **-2*log(L):** 75.1 | **df. resid:** 15 | |
| **Dispersion parameter for nbinom2 family ():** 2.27 | | | | | | |
| **Conditional Model:** | | ***Estimate*** | ***SE*** | ***z-value*** | ***Pr(>\|z\|)*** | |
| **(Intercept)** | | -0.6810 | 0.5495 | -1.239 | 0.215 |  |
| **Treatment** tunicamycin | | 2.7167 | 0.5992 | 4.534 | 5.78E-06 | *** |
| **Signif. codes:** 0 ‘***’ 0.001 ‘**’ 0.01 ‘*’ 0.05 ‘.’ 0.1 ‘ ’ 1 | | | | | | |

**b) Excretory Cell Total Area—Estimated Model Means**

| ***Treatment*** | ***response*** | ***SE*** | ***df*** | ***asymp.LCL*** | ***asymp.UCL*** |
| --- | --- | --- | --- | --- | --- |
| vehicle | 0.50612502 | 0.27811774 | Inf | 0.17239279 | 1.48592365 |
| tunicamycin | 7.65809990 | 1.82902272 | Inf | 4.79539359 | 12.2297561 |

**c) Excretory Cell Total Area —Contrasts**

| ***contrast*** | ***ratio*** | ***SE*** | ***df*** | ***null*** | ***z.ratio*** | ***p.value*** |  |
| --- | --- | --- | --- | --- | --- | --- | --- |
| vehicle / tunicamycin | 0.066090 | 0.039598 | Inf | 1 | - 4.5342 | 5.7818E-06 | *** |
| **Signif. codes:** 0 ‘***’ 0.001 ‘**’ 0.01 ‘*’ 0.05 ‘.’ 0.1 ‘ ’ 1 | | | | | | | |

**Online Resource 7 (Accompanies Fig. 2h)** Statistical analysis for differences in the number of Proteostat puncta in the excretory cell due to tunicamycin treatment. a) Raw data were fitted to a generalized linear model. The R package, ‘emmeans’ was used to obtain estimated marginal means on the response scale (b) and pairwise contrasts (c) with Tukey corrected p-values. Raw data are provided as a Source Data file

**a) Excretory Cell Count—Model Statistics**

| **Family:** nbinom2 (log) | | | | | | |
| --- | --- | --- | --- | --- | --- | --- |
| **Formula:** Count ~ Treatment | | | | | | |
| **AIC:** 121.1 | **BIC:** 123.8 | **logLik:** -57.6 | | **-2*log(L):** 115.1 | **df. resid:** 15 | |
| **Dispersion parameter for nbinom2 family ():** 1.96 | | | | | | |
| **Conditional Model:** | | ***Estimate*** | ***SE*** | ***z-value*** | ***Pr(>\|z\|)*** | |
| **(Intercept)** | | 1.1787 | 0.3200 | 3.683 | 0.00023 | *** |
| **Treatment** tunicamycin | | 1.7969 | 0.3983 | 4.511 | 6.45E-06 | *** |
| **Signif. codes:** 0 ‘***’ 0.001 ‘**’ 0.01 ‘*’ 0.05 ‘.’ 0.1 ‘ ’ 1 | | | | | | |

**b) Excretory Cell Count—Estimated Model Means**

| ***Treatment*** | ***response*** | ***SE*** | ***df*** | ***asymp.LCL*** | ***asymp.UCL*** |
| --- | --- | --- | --- | --- | --- |
| vehicle | 3.25000065 | 1.04000075 | Inf | 1.73579651 | 6.08510509 |
| tunicamycin | 19.6000005 | 4.64866738 | Inf | 12.3131945 | 31.1990541 |

**c) Excretory Cell Count—Contrasts**

| ***contrast*** | ***ratio*** | ***SE*** | ***df*** | ***null*** | ***z.ratio*** | ***p.value*** |  |
| --- | --- | --- | --- | --- | --- | --- | --- |
| vehicle / tunicamycin | 0.165816 | 0.0660467 | Inf | 1 | -4.51121 | 6.445e-06 | *** |
| **Signif. codes:** 0 ‘***’ 0.001 ‘**’ 0.01 ‘*’ 0.05 ‘.’ 0.1 ‘ ’ 1 | | | | | | | |

**Online Resource 8 (Accompanies Fig. 2i)** Statistical analysis for differences in the integrated density of Proteostat puncta in the excretory cell due to tunicamycin treatment. a) Raw data were fitted to a generalized linear model. The R package, ‘emmeans’ was used to obtain estimated marginal means on the response scale (b) and pairwise contrasts (c) with Tukey corrected p-values. Raw data are provided as a Source Data file

**a) Excretory Cell Integrated Density—Model Statistics**

| **Family:** nbinom2 (log) | | | | | | |
| --- | --- | --- | --- | --- | --- | --- |
| **Formula:** IntDen ~ Treatment | | | | | | |
| **AIC:** 165.6 | **BIC:** 168.3 | **logLik:** -79.8 | | **-2*log(L):** 159.6 | **df. resid:** 15 | |
| **Dispersion parameter for nbinom2 family ():**1.18 | | | | | | |
| **Conditional Model:** | | ***Estimate*** | ***SE*** | ***z-value*** | ***Pr(>\|z\|)*** | |
| **(Intercept)** | | 2.6480 | 0.3389 | 7.813 | 5.6E-15 | *** |
| **Treatment** tunicamycin | | 1.3863 | 0.4489 | 3.088 | 0.00201 | ** |
| **Signif. codes:** 0 ‘***’ 0.001 ‘**’ 0.01 ‘*’ 0.05 ‘.’ 0.1 ‘ ’ 1 | | | | | | |

**b) Excretory Cell Integrated Density—Estimated Model Means**

| ***Treatment*** | ***response*** | ***SE*** | ***df*** | ***asymp.LCL*** | ***asymp.UCL*** |
| --- | --- | --- | --- | --- | --- |
| vehicle | 14.1251248 | 4.78751430 | Inf | 7.26924635 | 27.4470201 |
| tunicamycin | 56.4986014 | 16.6256194 | Inf | 31.7364243 | 100.581336 |

**c) Excretory Cell Integrated Density—Contrasts**

| ***contrast*** | ***ratio*** | ***SE*** | ***df*** | ***null*** | ***z.ratio*** | ***p.value*** |  |
| --- | --- | --- | --- | --- | --- | --- | --- |
| vehicle / tunicamycin | 0.2500083 | 0.112217 | Inf | 1 | -3.0884 | 0.002012 | ** |
| **Signif. codes:** 0 ‘***’ 0.001 ‘**’ 0.01 ‘*’ 0.05 ‘.’ 0.1 ‘ ’ 1 | | | | | | | |

**Online Resource 9 (Accompanies Fig. 3g)** Statistical analysis for differences in the total area occupied by Proteostat puncta in the superficial head tissues due to hypoxia treatment. a) Raw data were fitted to a generalized linear model. The R package, ‘emmeans’ was used to obtain estimated marginal means on the response scale (b) and pairwise contrasts (c) with Tukey corrected p-values. Raw data are provided as a Source Data file

**a) Superficial Head Total Area—Model Statistics**

| **Family:** nbinom2 ( log ) | | | | | |
| --- | --- | --- | --- | --- | --- |
| **Formula:** Total Area ~ Treatment | | | | | |
| **AIC:** 90.4 | **BIC:** 92.9 | **logLik:** -42.2 | **-2*log(L):** 84.4 | **df.resid:** 14 | |
| **Dispersion parameter for nbinom2 family ():** 0.871 | | | | | |
| **Conditional model:** | **Estimate** | **Std. Error** | **z value** | **Pr(>\|z\|)** | |
| **(Intercept)** | -0.1213 | 0.5031 | -0.241 | 0.809 |  |
| **Treatment Hypoxia** | 2.9039 | 0.6359 | 4.567 | 4.96e-06 | *** |
| **Signif. codes:** 0 ‘***’ 0.001 ‘**’ 0.01 ‘*’ 0.05 ‘.’ 0.1 ‘ ’ 1 | | | | | |

**b) Superficial Head Total Area—Estimated Model Means**

| **Treatment** | **response** | **SE** | **df** | **asymp.LCL** | **asymp.UCL** |
| --- | --- | --- | --- | --- | --- |
| **Control** | 0.885777761 | 0.445600117 | Inf | 0.330459771 | 2.374274606 |
| **Hypoxia** | 16.16187516 | 6.286934021 | Inf | 7.540084497 | 34.64234503 |

**c) Superficial Head Total Area—Contrasts**

| **contrast** | **ratio** | **SE** | **df** | **null** | **z.ratio** | **p.value** |  |
| --- | --- | --- | --- | --- | --- | --- | --- |
| **Control / Hypoxia** | 0.054806621 | 0.034852422 | Inf | 1 | -4.566551226 | 4.96E-06 | *** |
| **Signif. codes:** 0 ‘***’ 0.001 ‘**’ 0.01 ‘*’ 0.05 ‘.’ 0.1 ‘ ’ 1 | | | | | | | |

**Online Resource 10 (Accompanies Fig. 3j)** Statistical analysis for differences in the number of Proteostat puncta in the superficial head tissues due to hypoxia treatment. a) Raw data were fitted to a generalized linear model. The R package, ‘emmeans’ was used to obtain estimated marginal means on the response scale (b) and pairwise contrasts (c) with Tukey corrected p-values. Raw data are provided as a Source Data file

**a) Superficial Head Count—Model Statistics**

| **Family:** nbinom2 ( log ) | | | | | |
| --- | --- | --- | --- | --- | --- |
| **Formula:** Count ~ Treatment | | | | | |
| **AIC:** 154.5 | **BIC:** 157.0 | **logLik:** -74.3 | **-2*log(L):** 148.5 | **df.resid:** 14 | |
| **Dispersion parameter for nbinom2 family ():** 0.567 | | | | | |
| **Conditional model:** | **Estimate** | **Std. Error** | **z value** | **Pr(>\|z\|)** | |
| **(Intercept)** | 2.5650 | 0.4524 | 5.670 | 1.43e-08 | *** |
| **Treatment Hypoxia** | 1.8372 | 0.6533 | 2.812 | 0.00492 | ** |
| **Signif. codes:** 0 ‘***’ 0.001 ‘**’ 0.01 ‘*’ 0.05 ‘.’ 0.1 ‘ ’ 1 | | | | | |

**b) Superficial Head Count—Estimated Model Means**

| **Treatment** | **response** | **SE** | **df** | **asymp.LCL** | **asymp.UCL** |
| --- | --- | --- | --- | --- | --- |
| **Control** | 13.00001475 | 5.880648496 | Inf | 5.356693709 | 31.54938337 |
| **Hypoxia** | 81.6250648 | 38.46967864 | Inf | 32.40816845 | 205.5855521 |

**c) Superficial Head Count—Contrasts**

| **contrast** | **ratio** | **SE** | **df** | **null** | **z.ratio** | **p.value** |  |
| --- | --- | --- | --- | --- | --- | --- | --- |
| **Control / Hypoxia** | 0.159264985 | 0.104041383 | Inf | 1 | -2.812336539 | 0.004918301 | ** |
| **Signif. codes:** 0 ‘***’ 0.001 ‘**’ 0.01 ‘*’ 0.05 ‘.’ 0.1 ‘ ’ 1 | | | | | | | |

**Online Resource 11 (Accompanies Fig. 3m)** Statistical analysis for differences in the integrated density of Proteostat puncta in the superficial head tissues due to hypoxia treatment. a) Raw data were fitted to a generalized linear model. The R package, ‘emmeans’ was used to obtain estimated marginal means on the response scale (b) and pairwise contrasts (c) with Tukey corrected p-values. Raw data are provided as a Source Data file

**a) Superficial Head Integrated Density—Model Statistics**

| **Family:** nbinom2 ( log ) | | | | | |
| --- | --- | --- | --- | --- | --- |
| **Formula:** IntDen ~ Treatment | | | | | |
| **AIC:** 131.7 | **BIC:** 134.2 | **logLik:** -62.8 | **-2*log(L):** 125.7 | **df.resid:** 14 | |
| **Dispersion parameter for nbinom2 family ():** 1.68 | | | | | |
| **Conditional model:** | **Estimate** | **Std. Error** | **z value** | **Pr(>\|z\|)** | |
| **(Intercept)** | 2.0010 | 0.2848 | 7.027 | 2.12e-12 | *** |
| **Treatment Hypoxia** | 1.4611 | 0.3992 | 3.660 | 0.000252 | *** |
| **Signif. codes:** 0 ‘***’ 0.001 ‘**’ 0.01 ‘*’ 0.05 ‘.’ 0.1 ‘ ’ 1 | | | | | |

**b) Superficial Head Integrated Density—Estimated Model Means**

| **Treatment** | **response** | **SE** | **df** | **asymp.LCL** | **asymp.UCL** |
| --- | --- | --- | --- | --- | --- |
| **Control** | 7.396223566 | 2.106221689 | Inf | 4.232660837 | 12.92428691 |
| **Hypoxia** | 31.88313578 | 8.918772233 | Inf | 18.42687459 | 55.165858 |

**c) Superficial Head Integrated Density—Contrasts**

| **contrast** | **ratio** | **SE** | **df** | **null** | **z.ratio** | **p.value** |  |
| --- | --- | --- | --- | --- | --- | --- | --- |
| **Control / Hypoxia** | 0.231979176 | 0.092601414 | Inf | 1 | -3.660274046 | 0.000251946 | *** |
| **Signif. codes:** 0 ‘***’ 0.001 ‘**’ 0.01 ‘*’ 0.05 ‘.’ 0.1 ‘ ’ 1 | | | | | | | |

**Online Resource 12 (Accompanies Fig. 3h)** Statistical analysis for differences in the total area occupied by Proteostat puncta in the deep head tissues due to hypoxia treatment. a) Raw data were fitted to a generalized linear model. The R package, ‘emmeans’ was used to obtain estimated marginal means on the response scale (b) and pairwise contrasts (c) with Tukey corrected p-values. Raw data are provided as a Source Data file

**a) Deep Head Total Area—Model Statistics**

| **Family:** nbinom2 ( log ) | | | | | |
| --- | --- | --- | --- | --- | --- |
| **Formula:** TotalArea ~ Treatment | | | | | |
| **AIC:** 85.9 | **BIC:** 88.9 | **logLik:** -39.9 | **-2*log(L):** 79.9 | **df.resid:** 17 | |
| **Dispersion parameter for nbinom2 family ():** 5.59 | | | | | |
| **Conditional model:** | **Estimate** | **Std. Error** | **z value** | **Pr(>\|z\|)** | |
| **(Intercept)** | 0.2725 | 0.2924 | 0.932 | 0.351 |  |
| **Treatment Hypoxia** | 1.5468 | 0.3513 | 4.404 | 1.07e-05 | *** |
| **Signif. codes:** 0 ‘***’ 0.001 ‘**’ 0.01 ‘*’ 0.05 ‘.’ 0.1 ‘ ’ 1 | | | | | |

**b) Deep Head Total Area—Estimated Model Means**

| **Treatment** | **response** | **SE** | **df** | **asymp.LCL** | **asymp.UCL** |
| --- | --- | --- | --- | --- | --- |
| **Control** | 1.313181901 | 0.383968227 | Inf | 0.740350801 | 2.3292292 |
| **Hypoxia** | 6.167444202 | 1.200634163 | Inf | 4.211136975 | 9.032564889 |

**c) Deep Head Total Area—Contrasts**

| **contrast** | **ratio** | **SE** | **df** | **null** | **z.ratio** | **p.value** |  |
| --- | --- | --- | --- | --- | --- | --- | --- |
| **Control / Hypoxia** | 0.21292157 | 0.074793547 | Inf | 1 | -4.403505178 | 1.07E-05 | *** |
| **Signif. codes:** 0 ‘***’ 0.001 ‘**’ 0.01 ‘*’ 0.05 ‘.’ 0.1 ‘ ’ 1 | | | | | | | |

**Online Resource 13 (Accompanies Fig. 3k)** Statistical analysis for differences in the number of Proteostat puncta in the deep head tissues due to hypoxia treatment. a) Raw data were fitted to a generalized linear model. The R package, ‘emmeans’ was used to obtain estimated marginal means on the response scale (b) and pairwise contrasts (c) with Tukey corrected p-values. Raw data are provided as a Source Data file

**a) Deep Head Count—Model Statistics**

| **Family:** nbinom2 ( log ) | | | | | |
| --- | --- | --- | --- | --- | --- |
| **Formula:** Count ~ Treatment | | | | | |
| **AIC**: 184.5 | **BIC:** 187.5 | **logLik:** -89.3 | **-2*log(L):** 178.5 | **df.resid:** 17 | |
| **Dispersion parameter for nbinom2 family ():** 1.44 | | | | | |
| **Conditional model:** | **Estimate** | **Std. Error** | **z value** | **Pr(>\|z\|)** | |
| **(Intercept)** | 3.0138 | 0.2601 | 11.587 | < 2e-16 | *** |
| **Treatment Hypoxia** | 1.0136 | 0.3832 | 2.645 | 0.00817 | ** |
| **Signif. codes:** 0 ‘***’ 0.001 ‘**’ 0.01 ‘*’ 0.05 ‘.’ 0.1 ‘ ’ 1 | | | | | |

**b) Deep Head Count— Estimated Model Means**

| **Treatment** | **response** | **SE** | **df** | **asymp.LCL** | **asymp.UCL** |
| --- | --- | --- | --- | --- | --- |
| **Control** | 20.3636353 | 5.296747438 | Inf | 12.23068707 | 33.90468911 |
| **Hypoxia** | 56.11111889 | 15.79254695 | Inf | 32.32043179 | 97.41384902 |

**c) Deep Head Count— Contrasts**

| **contrast** | **ratio** | **SE** | **df** | **null** | **z.ratio** | **p.value** |  |
| --- | --- | --- | --- | --- | --- | --- | --- |
| **Control / Hypoxia** | 0.362916222 | 0.139083152 | Inf | 1 | -2.64479058 | 0.008174149 | ** |
| **Signif. codes:** 0 ‘***’ 0.001 ‘**’ 0.01 ‘*’ 0.05 ‘.’ 0.1 ‘ ’ 1 | | | | | | | |

**Online Resource 14 (Accompanies Fig. 3n)** Statistical analysis for differences in the integrated density of Proteostat puncta in the deep head tissues due to hypoxia treatment. a) Raw data were fitted to a generalized linear model. The R package, ‘emmeans’ was used to obtain estimated marginal means on the response scale (b) and pairwise contrasts (c) with Tukey corrected p-values. Raw data are provided as a Source Data file

**a) Deep Head Integrated Density—Model Statistics**

| **Family:** nbinom2 ( log ) | | | | | | | |
| --- | --- | --- | --- | --- | --- | --- | --- |
| **Formula:** IntDen ~ Treatment | | | | | | | |
| **AIC:** 123.5 | **BIC:** 126.3 | | **logLik:** -58.7 | **-2*log(L):** 117.5 | | **df.resid:** 16 | |
| **Dispersion parameter for nbinom2 family ():** 7.34 | | | | | | | |
| **Conditional model:** | | **Estimate** | **Std. Error** | **z value** | **Pr(>\|z\|)** | | |
| **(Intercept)** | | 2.0797 | 0.1616 | 12.867 | < 2e-16 | | *** |
| **Treatment Hypoxia** | | 0.6895 | 0.2196 | 3.139 | 0.00169 | | ** |
| **Signif. codes:** 0 ‘***’ 0.001 ‘**’ 0.01 ‘*’ 0.05 ‘.’ 0.1 ‘ ’ 1 | | | | | | | |

**b) Deep Head Integrated Density—Estimated Model Means**

| **Treatment** | **response** | **SE** | **df** | **asymp.LCL** | **asymp.UCL** |
| --- | --- | --- | --- | --- | --- |
| **Control** | 8.002199979 | 1.293394828 | Inf | 5.829479445 | 10.98472087 |
| **Hypoxia** | 15.94544468 | 2.370999725 | Inf | 11.91427604 | 21.34055021 |

**c) Deep Head Integrated Density—Contrasts**

| **contrast** | **ratio** | **SE** | **df** | **null** | **z.ratio** | **p.value** |  |
| --- | --- | --- | --- | --- | --- | --- | --- |
| **Control / Hypoxia** | 0.501848656 | 0.110217524 | Inf | 1 | -3.13927312 | 0.001693675 | ** |
| **Signif. codes:** 0 ‘***’ 0.001 ‘**’ 0.01 ‘*’ 0.05 ‘.’ 0.1 ‘ ’ 1 | | | | | | | |

**Online Resource 15 (Accompanies Fig. 3i)** Statistical analysis for differences in the total area occupied by Proteostat puncta in the excretory cell due to hypoxia treatment. a) Raw data were fitted to a generalized linear model. The R package, ‘emmeans’ was used to obtain estimated marginal means on the response scale (b) and pairwise contrasts (c) with Tukey corrected p-values. Raw data are provided as a Source Data file

**a) Excretory Cell Total Area—Model Statistics**

| **Family:** nbinom2 ( log ) | | | | | | |
| --- | --- | --- | --- | --- | --- | --- |
| **Formula:** `Total Area` ~ Treatment | | | | | | |
| **AIC:** 64.8 | **BIC:** 67.9 | **logLik:** -29.4 | **-2*log(L):** 58.8 | | **df.resid:** 18 | |
| **Dispersion parameter for nbinom2 family ():** 2.95 | | | | | | |
| **Conditional model:** | **Estimate** | **Std. Error** | | **z value** | **Pr(>\|z\|)** | |
| **(Intercept)** | -1.6484 | 0.7095 | | -2.323 | 0.0202 | * |
| **Treatment Hypoxia** | 3.0121 | 0.7502 | | 4.015 | 5.94e-05 | *** |
| **Signif. codes:** 0 ‘***’ 0.001 ‘**’ 0.01 ‘*’ 0.05 ‘.’ 0.1 ‘ ’ 1 | | | | | | |

**b) Excretory Cell Total Area—Estimated Model Means**

| **Treatment** | **response** | **SE** | **df** | **asymp.LCL** | **asymp.UCL** |
| --- | --- | --- | --- | --- | --- |
| **Control** | 0.192363639 | 0.136478797 | Inf | 0.047886123 | 0.772745165 |
| **Hypoxia** | 3.910699971 | 0.953324366 | Inf | 2.425237422 | 6.306011164 |

**c) Excretory Cell Total Area—Contrasts**

| **contrast** | **ratio** | **SE** | **df** | **null** | **z.ratio** | **p.value** |  |
| --- | --- | --- | --- | --- | --- | --- | --- |
| **Control / Hypoxia** | 0.049189056 | 0.036901368 | Inf | 1 | -4.015070007 | 5.94E-05 | *** |
| **Signif. codes:** 0 ‘***’ 0.001 ‘**’ 0.01 ‘*’ 0.05 ‘.’ 0.1 ‘ ’ 1 | | | | | | | |

**Online Resource 16 (Accompanies Fig 3l)** Statistical analysis for differences in the number of Proteostat puncta in the excretory cell due to hypoxia treatment. a) Raw data were fitted to a generalized linear model. The R package, ‘emmeans’ was used to obtain estimated marginal means on the response scale (b) and pairwise contrasts (c) with Tukey corrected p-values. Raw data are provided as a Source Data file

**a) Excretory Cell Count—Model Statistics**

| **Family:** nbinom2 ( log ) | | | | | |
| --- | --- | --- | --- | --- | --- |
| **Formula:** Count ~ Treatment | | | | | |
| **AIC:** 125.3 | **BIC:** 128.5 | **logLik:** -59.7 | **-2*log(L):** 119.3 | **df.resid:** 18 | |
| **Dispersion parameter for nbinom2 family ():** 2.15 | | | | | |
| **Conditional model:** | **Estimate** | **Std. Error** | **z value** | **Pr(>\|z\|)** | |
| **(Intercept)** | 0.7376 | 0.2928 | 2.519 | 0.0118 | * |
| **Treatment Hypoxia** | 2.2068 | 0.3707 | 5.953 | 2.63e-09 | *** |
| **Signif. codes:** 0 ‘***’ 0.001 ‘**’ 0.01 ‘*’ 0.05 ‘.’ 0.1 ‘ ’ 1 | | | | | |

**b) Excretory Cell Count—Estimated Model Means**

| **Treatment** | **response** | **SE** | **df** | **asymp.LCL** | **asymp.UCL** |
| --- | --- | --- | --- | --- | --- |
| **Control** | 2.090909016 | 0.612146813 | Inf | 1.17796495 | 3.711401186 |
| **Hypoxia** | 18.9999968 | 4.320989335 | Inf | 12.16670063 | 29.67114006 |

**c) Excretory Cell Count—Contrasts**

| **contrast** | **ratio** | **SE** | **df** | **null** | **z.ratio** | **p.value** |  |
| --- | --- | --- | --- | --- | --- | --- | --- |
| **Control / Hypoxia** | 0.110047861 | 0.040796741 | Inf | 1 | -5.952877749 | 2.63E-09 | *** |
| **Signif. codes:** 0 ‘***’ 0.001 ‘**’ 0.01 ‘*’ 0.05 ‘.’ 0.1 ‘ ’ 1 | | | | | | | |

**Online Resource 17 (Accompanies Fig. 3o)** Statistical analysis for differences in the integrated density of Proteostat puncta in the excretory cell due to hypoxia treatment. a) Raw data were fitted to a generalized linear model. The R package, ‘emmeans’ was used to obtain estimated marginal means on the response scale (b) and pairwise contrasts (c) with Tukey corrected p-values. Raw data are provided as a Source Data file

**a) Excretory Cell Integrated Density—Model Statistics**

| **Family:** nbinom2 ( log ) | | | | | |
| --- | --- | --- | --- | --- | --- |
| **Formula:** IntDen ~ Treatment | | | | | |
| **AIC:** 161.0 | **BIC:** 164.1 | **logLik:** -77.5 | **-2*log(L):** 155.0 | **df.resid:** 18 | |
| **Dispersion parameter for nbinom2 family ():** 1.38 | | | | | |
| **Conditional model:** | **Estimate** | **Std. Error** | **z value** | **Pr(>\|z\|)** | |
| **(Intercept)** | 1.9508 | 0.2807 | 6.950 | 3.66e-12 | *** |
| **Treatment Hypoxia** | 1.4968 | 0.3930 | 3.809 | 0.00014 | *** |
| **Signif. codes:** 0 ‘***’ 0.001 ‘**’ 0.01 ‘*’ 0.05 ‘.’ 0.1 ‘ ’ 1 | | | | | |

**b) Excretory Cell Integrated Density—Estimated Model Means**

| **Treatment** | **response** | **SE** | **df** | **asymp.LCL** | **asymp.UCL** |
| --- | --- | --- | --- | --- | --- |
| **Control** | 7.034448149 | 1.974564976 | Inf | 4.057871765 | 12.19443679 |
| **Hypoxia** | 31.42550568 | 8.642744121 | Inf | 18.33082772 | 53.87440339 |

**c) Excretory Cell Integrated Density—Contrasts**

| **contrast** | **ratio** | **SE** | **df** | **null** | **z.ratio** | **p.value** |  |
| --- | --- | --- | --- | --- | --- | --- | --- |
| **Control / Hypoxia** | 0.223845186 | 0.087965718 | Inf | 1 | -3.808888463 | 0.000139593 | *** |
| **Signif. codes:** 0 ‘***’ 0.001 ‘**’ 0.01 ‘*’ 0.05 ‘.’ 0.1 ‘ ’ 1 | | | | | | | |

**Online Resource 18 (Accompanies Fig. 4e)** Statistical analysis for differences in the integrated density of Proteostat puncta in the excretory cell due to genotype and tunicamycin treatment. a) Raw data were fitted to a generalized linear model. The R package, ‘emmeans’ was used to obtain estimated marginal means on the response scale (b) and pairwise contrasts (c) with Tukey corrected p-values. Raw data are provided as a Source Data file

**a) Excretory Cell Integrated Density—Model Statistics**

| **Family:** gaussian ( identity ) | | | | | | | |
| --- | --- | --- | --- | --- | --- | --- | --- |
| **Formula:** IntDen ~ Genotype * Treatment | | | | | | | |
| **Dispersion:** ~Treatment | | | | | | | |
| **AIC:** 323.7 | **BIC:** 333.5 | | **logLik:** -155.8 | | -**2*log(L):** 311.7 | **df.resid:** 32 | |
| **Conditional model:** | | **Estimate** | | **Std. Error** | **z value** | **Pr(>\|z\|)** | |
| (Intercept) | | 14.125 | | 3.912 | 3.610 | 0.000306 | *** |
| Genotype *hsp-4(gk514)* | | -8.814 | | 5.249 | -1.679 | 0.093126 | . |
| Treatment Tunicamycin | | 42.373 | | 7.108 | 5.962 | 2.5e-09 | *** |
| Genotype *hsp-4(gk514)*: Treatment Tunicamycin | | -31.590 | | 9.898 | -3.191 | 0.001415 | ** |
| **Signif. codes:** 0 ‘***’ 0.001 ‘**’ 0.01 ‘*’ 0.05 ‘.’ 0.1 ‘ ’ 1 | | | | | | | |
| **Dispersion model:** | | **Estimate** | | **Std. Error** | **z ratio** | **Pr(>\|z\|)** | |
| (Intercept) | | 2.4039 | | 0.1667 | 14.423 | <2e-16 | *** |
| Treatment Tunicamycin | | 0.5281 | | 0.2297 | 2.299 | 0.0215 | * |
| **Signif. codes:** 0 ‘***’ 0.001 ‘**’ 0.01 ‘*’ 0.05 ‘.’ 0.1 ‘ ’ 1 | | | | | | | |

**b) Excretory Cell Integrated Density—Estimated Model Means**

| **Genotype** | **Treatment** | **emmean** | **SE** | **df** | **asymp.LCL** | **asymp.UCL** |
| --- | --- | --- | --- | --- | --- | --- |
| wildtype | vehicle | 14.12512 | 3.912480 | Inf | 6.45680 | 21.79344 |
| *hsp-4(gk514)* | vehicle | 5.311103 | 3.499429 | Inf | -1.547651 | 12.16985 |
| wildtype | tunicamycin | 56.49858 | 5.93390 | Inf | 44.86834 | 68.12883 |
| *hsp-4(gk514)* | tunicamycin | 16.09430 | 5.93390 | Inf | 4.464061 | 27.724554 |

**c) Excretory Cell Integrated Density—Contrasts**

| **contrast** | **estimate** | **SE** | **df** | **z.ratio** | **p.value** |  |
| --- | --- | --- | --- | --- | --- | --- |
| wildtype vehicle /  *hsp-4(gk514)* vehicle | 8.814020 | 5.249143 | Inf | 1.679134 | 0.334717 |  |
| wildtype vehicle /  wildtype tunicamycin | -42.37346 | 7.107656 | Inf | -5.961664 | 1.50E-08 | *** |
| wildtype vehicle /  *hsp-4(gk514)* tunicamycin | -1.96918 | 7.107656 | Inf | -0.27705 | 0.992577 |  |
| *hsp-4(gk514)* vehicle / wildtype tunicamycin | -51.18748 | 6.888923 | Inf | -7.430403 | 6.76E-13 | *** |
| *hsp-4(gk514)* vehicle /  *hsp-4(gk514)* tunicamycin | -10.7832 | 6.888923 | Inf | -1.565295 | 0.398603 |  |
| wildtype tunicamycin /  *hsp-4(gk514)* tunicamycin | 40.40428 | 8.391813 | Inf | 4.814725 | 8.77E-06 | *** |
| **Signif. codes:** 0 ‘***’ 0.001 ‘**’ 0.01 ‘*’ 0.05 ‘.’ 0.1 ‘ ’ 1 | | | | | | |

**Online Resource 19 (Accompanies Fig. 5g)** Statistical analysis for differences in the integrated density of Proteostat puncta in the tissues near the rectum due to genotype and hypoxia treatment. a) Raw data were fitted to a generalized linear model. The R package, ‘emmeans’ was used to obtain estimated marginal means on the response scale (b) and pairwise contrasts (c) with Tukey corrected p-values. Raw data are provided as a Source Data file

**a) Rectal Tissues Integrated Density—Model Statistics**

| **Family:** gaussian ( identity ) | | | | | | | |
| --- | --- | --- | --- | --- | --- | --- | --- |
| **Formula:** IntDen ~ Strain * Treatment | | | | | | | |
| **Dispersion:** ~Treatment + Strain | | | | | | | |
| **AIC:** 830.8 | **BIC:** 843.2 | | **logLik:** -408.4 | | -**2*log(L):** 816.8 | **df.resid:** 36 | |
| **Dispersion estimate for gaussian family (sigma^2): 1.04e07** | | | | | | | |
| **Conditional model:** | | **Estimate** | | **Std. Error** | **z value** | **Pr(>\|z\|)** | |
| (Intercept) | | 7437 | | 1317 | 5.645 | 1.65e-08 | *** |
| Genotype *hsp-12.6(ok1077)* | | 7370 | | 1701 | 4.334 | 1.47e-05 | *** |
| Genotype *hsp-16.2(gk249)* | | 2121 | | 1863 | 1.138 | 0.25501 |  |
| Treatment Hypoxia | | 5592 | | 1863 | 3.002 | 0.00269 | ** |
| Genotype *hsp-12.6(ok1077)*: Treatment Hypoxia | | -2273 | | 2381 | -0.955 | 0.33974 |  |
| Genotype *hsp-16.2(gk249)*: Treatment Hypoxia | | -2268 | | 2635 | -0.861 | 0.38933 |  |
| **Signif. codes:** 0 ‘***’ 0.001 ‘**’ 0.01 ‘*’ 0.05 ‘.’ 0.1 ‘ ’ 1 | | | | | | | |
| **Dispersion model:** | | **Estimate** | | **Std. Error** | **z value** | **Pr(>\|z\|)** | |
| (Intercept) | | -0.7372 | | 0.6662 | -1.107 | 0.268 |  |
| Treatment Hypoxia | | 0.8345 | | 0.6245 | 1.336 | 0.181 |  |
| Genotype *hsp-12.6(ok1077)* | | 3.2255 | | 0.8209 | 3.929 | 8.53E-05 | *** |
| Genotype *hsp-16.2(gk249)* | | 1.1337 | | 0.7087 | 1.600 | 0.110 |  |
| **Signif. codes:** 0 ‘***’ 0.001 ‘**’ 0.01 ‘*’ 0.05 ‘.’ 0.1 ‘ ’ 1 | | | | | | | |

**b) Rectal Tissues Integrated Density—Estimated Model Means**

| **Genotype** | **Treatment** | **emmean** | **SE** | **df** | **lower.CL** | **upper.CL** |
| --- | --- | --- | --- | --- | --- | --- |
| reference | Control | 7436.9582 | 1317.397 | 36 | 4765.15181 | 10108.7646 |
| *hsp-12.6(ok1077)* | Control | 14807.415 | 1075.650 | 36 | 12625.8948 | 16988.9364 |
| *hsp-16.2(gk249)* | Control | 9557.6300 | 1317.397 | 36 | 6885.82365 | 12229.4365 |
| reference | Hypoxia | 13029.144 | 1317.397 | 36 | 10357.3382 | 15700.9511 |
| *hsp-12.6(ok1077)* | Hypoxia | 18126.471 | 1020.451 | 36 | 16056.8993 | 20196.044 |
| *hsp-16.2(gk249)* | Hypoxia | 12881.692 | 1317.397 | 36 | 10209.8857 | 15553.4986 |

**c) Rectal Tissues Integrated Density—Contrasts**

| **contrast** | **estimate** | **SE** | **df** | **t.ratio** | **p.value** |  |
| --- | --- | --- | --- | --- | --- | --- |
| wildtype Control /  (*hsp-12.6(ok1077)* Control*)* | -7370.4573 | 1700.7531 | 36 | -4.3336432 | 0.001466 | ** |
| wildtype Control /  (*hsp-16.2(gk249)* Control*)* | -2120.6718 | 1863.0817 | 36 | -1.1382602 | 0.862067 |  |
| wildtype Control /  wildtype Hypoxia | -5592.1864 | 1863.0817 | 36 | -3.0015787 | 0.050828 | * |
| wildtype Control /  (*hsp-12.6(ok1077)* Hypoxia) | -10689.513 | 1666.3909 | 36 | -6.4147692 | 2.78E-06 | *** |
| wildtype Control /  (*hsp-16.2(gk249)* Hypoxia) | -5444.7339 | 1863.0817 | 36 | -2.9224343 | 0.061074 | . |
| (*hsp-12.6(ok1077)* Control)/  (*hsp-16.2(gk249)* Control) | 5249.7855 | 1700.7531 | 36 | 3.0867416 | 0.041531 | * |
| (*hsp-12.6(ok1077)* Control)/  wildtype Hypoxia | 1778.2709 | 1700.7531 | 36 | 1.0455785 | 0.899090 |  |
| (*hsp-12.6(ok1077)* Control)/  (*hsp-12.6(ok1077)* Hypoxia)/ | -3319.0561 | 1482.6822 | 36 | -2.2385485 | 0.245854 |  |
| (*hsp-12.6(ok1077)* Control)/  (*hsp-16.2(gk249)* Hypoxia) | 1925.7234 | 1700.7531 | 36 | 1.1322768 | 0.864650 |  |
| (*hsp-16.2(gk249)* Control)/  wildtype Hypoxia | -3471.5146 | 1863.0817 | 36 | -1.8633185 | 0.440210 |  |
| (*hsp-16.2(gk249)* Control)/  (*hsp-12.6(ok1077)* Hypoxia) | -8568.8416 | 1666.3909 | 36 | -5.1421556 | 0.000133 | *** |
| (*hsp-16.2(gk249)* Control)/  (*hsp-16.2(gk249)* Hypoxia)/ | -3324.0621 | 1863.0817 | 36 | -1.7841741 | 0.488180 |  |
| wildtype Hypoxia/  (*hsp-12.6(ok1077)* Hypoxia) | -5097.327 | 1666.3909 | 36 | -3.0589022 | 0.044387 | * |
| wildtype Hypoxia/  (*hsp-16.2(gk249)* Hypoxia) | 147.45249 | 1863.0817 | 36 | 0.0791444 | 0.999999 |  |
| (*hsp-12.6(ok1077)* Hypoxia)/  (*hsp-16.2(gk249)* Hypoxia) | 5244.7794 | 1666.3909 | 36 | 3.1473883 | 0.035871 | * |

**Online Resource 20 (Accompanies Fig. 5h)** Statistical analysis for differences in the total area occupied by Proteostat puncta in the tissues near the rectum due to genotype and hypoxia treatment. a) Raw data were fitted to a generalized linear model. The R package, ‘emmeans’ was used to obtain estimated marginal means on the response scale (b) and pairwise contrasts (c) with Tukey corrected p-values. Raw data are provided as a Source Data file

**a) Rectal Tissues Total Area—Model Statistics**

| **Family:** gaussian ( identity ) | | | | | | | |
| --- | --- | --- | --- | --- | --- | --- | --- |
| **Formula:** TotalArea ~ Strain * Treatment | | | | | | | |
| **Dispersion:** ~ Strain * Treatment | | | | | | | |
| **AIC:** 274.6 | **BIC:** 295.7 | | **logLik:** -125.3 | | -**2*log(L):** 250.6 | **df.resid:** 31 | |
| **Conditional model:** | | **Estimate** | | **Std. Error** | **z value** | **Pr(>\|z\|)** | |
| (Intercept) | | 0.3632 | | 0.3315 | 1.095 | 0.273323 |  |
| Genotype *hsp-12.6(ok1077)* | | 6.3356 | | 1.7612 | 3.597 | 0.000321 | *** |
| Genotype *hsp-16.2(gk249)* | | 1.7890 | | 1.4270 | 1.254 | 0.209941 |  |
| Treatment Hypoxia | | 9.4967 | | 3.5972 | 2.640 | 0.008290 | ** |
| Genotype *hsp-12.6(ok1077)*: Treatment Hypoxia | | -3.5568 | | 4.5771 | -0.777 | 0.437112 |  |
| Genotype *hsp-16.2(gk249)*: Treatment Hypoxia | | -2.7172 | | 4.5718 | -0.594 | 0.552284 |  |
| **Dispersion model:** | | **Estimate** | | **Std. Error** | **z ratio** | **Pr(>\|z\|)** |  |
| (Intercept) | | -0.2082 | | 0.2887 | -0.721 | 0.470819 |  |
| Genotype *hsp-12.6(ok1077)* | | 1.8547 | | 0.3727 | 4.977 | 6.46e-07 | *** |
| Genotype *hsp-16.2(gk249)* | | 1.4319 | | 0.4082 | 3.507 | 0.000453 | *** |
| Treatment Hypoxia | | 2.3799 | | 0.4082 | 5.830 | 5.55e-09 | *** |
| Genotype *hsp-12.6(ok1077)*: Treatment Hypoxia | | -2.0687 | | 0.5217 | -3.965 | 7.34e-05 | *** |
| Genotype *hsp-16.2(gk249)*: Treatment Hypoxia | | -1.8090 | | 0.5774 | -3.133 | 0.001729 | ** |
| **Signif. codes:** 0 ‘***’ 0.001 ‘**’ 0.01 ‘*’ 0.05 ‘.’ 0.1 ‘ ’ 1 | | | | | | | |

**b) Rectal Tissues Count—Estimated Model Means**

| **Genotype** | **Treatment** | **emmean** | **SE** | **df** | **asymp.LCL** | **asymp.UCL** |
| --- | --- | --- | --- | --- | --- | --- |
| wildtype | Control | 0.3631651 | 0.331523 | Inf | -0.2866092 | 1.01293945 |
| *hsp-12.6(ok1077)* | Control | 6.6987907 | 1.729714 | Inf | 3.30861268 | 10.0889688 |
| *hsp-16.2(gk249)* | Control | 2.1522100 | 1.387934 | Inf | -0.5680909 | 4.87251091 |
| wildtype | Hypoxia | 9.8598341 | 3.581879 | Inf | 2.83947998 | 16.8801884 |
| *hsp-12.6(ok1077)* | Hypoxia | 12.638672 | 2.240163 | Inf | 8.24803242 | 17.0293127 |
| *hsp-16.2(gk249)* | Hypoxia | 8.9316697 | 2.456671 | Inf | 4.11668215 | 13.7466573 |

**c) Rectal Tissues Count—Contrasts**

| **contrast** | **estimate** | **SE** | **df** | **z.ratio** | **p.value** |  |
| --- | --- | --- | --- | --- | --- | --- |
| wildtype Control /  (*hsp-12.6(ok1077)* Control*)* | -6.3356256 | 1.7611984 | Inf | -3.5973377 | 0.004345 | ** |
| wildtype Control /  (*hsp-16.2(gk249)* Control*)* | -1.7890449 | 1.4269789 | Inf | -1.253729 | 0.810064 |  |
| wildtype Control /  wildtype Hypoxia | -9.4966691 | 3.5971886 | Inf | -2.6400253 | 0.087829 | . |
| wildtype Control /  (*hsp-12.6(ok1077)* Hypoxia) | -12.275507 | 2.2645620 | Inf | -5.4206982 | 8.86E-07 | *** |
| wildtype Control /  (*hsp-16.2(gk249)* Hypoxia) | -8.5685046 | 2.4789397 | Inf | -3.4565198 | 0.007235 | ** |
| (*hsp-12.6(ok1077)* Control)/  (*hsp-16.2(gk249)* Control) | 4.5465807 | 2.2177180 | Inf | 2.0501166 | 0.313942 |  |
| (*hsp-12.6(ok1077)* Control)/  wildtype Hypoxia | -3.1610435 | 3.9776589 | Inf | -0.7946995 | 0.968509 |  |
| (*hsp-12.6(ok1077)* Control)/  (*hsp-12.6(ok1077)* Hypoxia)/ | -5.9398818 | 2.8302376 | Inf | -2.0987219 | 0.287809 |  |
| (*hsp-12.6(ok1077)* Control)/  (*hsp-16.2(gk249)* Hypoxia) | -2.232879 | 3.0045210 | Inf | -0.743173 | 0.976506 |  |
| (*hsp-16.2(gk249)* Control)/  wildtype Hypoxia | -7.7076242 | 3.8413825 | Inf | -2.0064714 | 0.338463 | . |
| (*hsp-16.2(gk249)* Control)/  (*hsp-12.6(ok1077)* Hypoxia) | -10.486463 | 2.6352788 | Inf | -3.9792611 | 0.000976 | *** |
| (*hsp-16.2(gk249)* Control)/  (*hsp-16.2(gk249)* Hypoxia)/ | -6.7794597 | 2.8216299 | Inf | -2.402675 | 0.155160 |  |
| wildtype Hypoxia/  (*hsp-12.6(ok1077)* Hypoxia) | -2.7788384 | 4.2247120 | Inf | -0.6577581 | 0.986394 |  |
| wildtype Hypoxia/  (*hsp-16.2(gk249)* Hypoxia) | 0.9281644 | 4.3433964 | Inf | 0.2136955 | 0.999939 |  |
| (*hsp-12.6(ok1077)* Hypoxia)/  (*hsp-16.2(gk249)* Hypoxia) | 3.7070028 | 3.3246906 | Inf | 1.1149918 | 0.875398 |  |

**Online Resource 21 (Accompanies Fig. 5i)** Statistical analysis for differences in the mean fluorescence of Proteostat in the posterior intestine due to genotype and hypoxia treatment. a) Raw data were fitted to a generalized linear model. The R package, ‘emmeans’ was used to obtain estimated marginal means on the response scale (b) and pairwise contrasts (c) with Tukey corrected p-values. Raw data are provided as a Source Data file.

**a) Posterior Intestine Mean Intensity—Model Statistics**

| **Family:** gaussian (identity) | | | | | | | |
| --- | --- | --- | --- | --- | --- | --- | --- |
| **Formula:** Mean ~ Strain * Treatment | | | | | | | |
| **AIC:** 372.7 | **BIC:** 388.6 | | **logLik:** -177.4 | | -**2*log(L):** 354.7 | **df.resid:** 34 | |
| **Conditional model:** | | **Estimate** | | **Std. Error** | **z value** | **Pr(>\|z\|)** | |
| (Intercept) | | 48.889 | | 5.108 | 9.571 | <2e-16 | *** |
| Genotype *hsp-12.6(ok1077)* | | 18.134 | | 8.730 | 2.077 | 0.0378 | * |
| Genotype *hsp-16.2(gk249)* | | -5.003 | | 6.611 | -0.757 | 0.4492 |  |
| Treatment Hypoxia | | 5.899 | | 7.224 | 0.817 | 0.4141 |  |
| Genotype *hsp-12.6(ok1077)*: Treatment Hypoxia | | 6.301 | | 12.141 | 0.519 | 0.6038 |  |
| Genotype *hsp-16.2(gk249)*: Treatment Hypoxia | | 1.310 | | 9.350 | 0.140 | 0.8886 |  |
| **Dispersion model:** | | **Estimate** | | **Std. Error** | **z value** | **Pr(>\|z\|)** |  |
| (Intercept) | | 2.5267 | | 0.2041 | 12.378 | <2e-16 | *** |
| Genotype *hsp-12.6(ok1077)* | | 0.5291 | | 0.2607 | 2.029 | 0.042 | * |
| Genotype *hsp-16.2(gk249)* | | -0.1964 | | 0.2887 | -0.680 | 0.4963 |  |
| **Signif. codes:** 0 ‘***’ 0.001 ‘**’ 0.01 ‘*’ 0.05 ‘.’ 0.1 ‘ ’ 1 | | | | | | | |

**b) Posterior Intestine Mean Intensity —Estimated Model Means**

| **Genotype** | **Treatment** | **emmean** | **SE** | **df** | **asymp.LCL** | **asymp.UCL** |
| --- | --- | --- | --- | --- | --- | --- |
| wildtype | Control | 48.888533 | 5.107979 | Inf | 38.8770777 | 58.8999884 |
| *hsp-12.6(ok1077)* | Control | 67.022656 | 7.079472 | Inf | 53.1471451 | 80.8981682 |
| *hsp-16.2(gk249)* | Control | 43.885168 | 4.197218 | Inf | 35.6587709 | 52.1115653 |
| wildtype | Hypoxia | 54.788007 | 5.107979 | Inf | 44.7765525 | 64.7994632 |
| *hsp-12.6(ok1077)* | Hypoxia | 79.222921 | 6.716177 | Inf | 66.0594552 | 92.3863873 |
| *hsp-16.2(gk249)* | Hypoxia | 51.094161 | 4.197218 | Inf | 42.867764 | 59.3205584 |

**c) Posterior Intestine Mean Intensity—Contrasts**

| **contrast** | **estimate** | **SE** | **df** | **z.ratio** | **p.value** |  |
| --- | --- | --- | --- | --- | --- | --- |
| wildtype Control /  (*hsp-12.6(ok1077)* Control*)* | -18.134124 | 8.7298559 | Inf | -2.0772535 | 0.299195 |  |
| wildtype Control /  (*hsp-16.2(gk249)* Control*)* | 5.0033649 | 6.6112098 | Inf | 0.7568002 | 0.974551 |  |
| wildtype Control /  wildtype Hypoxia | -5.8994748 | 7.2237735 | Inf | -0.8166749 | 0.964579 |  |
| wildtype Control /  (*hsp-12.6(ok1077)* Hypoxia) | -30.334388 | 8.4379199 | Inf | -3.5950078 | 0.004383 | ** |
| wildtype Control /  (*hsp-16.2(gk249)* Hypoxia) | -2.2056282 | 6.6112098 | Inf | -0.3336194 | 0.999455 |  |
| (*hsp-12.6(ok1077)* Control)/  (*hsp-16.2(gk249)* Control) | 23.137488 | 8.2301626 | Inf | 2.8113039 | 0.055665 | . |
| (*hsp-12.6(ok1077)* Control)/  wildtype Hypoxia | 12.234648 | 8.7298559 | Inf | 1.4014720 | 0.726222 |  |
| (*hsp-12.6(ok1077)* Control)/  (*hsp-12.6(ok1077)* Hypoxia)/ | -12.200265 | 9.7583797 | Inf | -1.2502347 | 0.811883 |  |
| (*hsp-12.6(ok1077)* Control)/  (*hsp-16.2(gk249)* Hypoxia) | 15.928495 | 8.2301626 | Inf | 1.9353804 | 0.380365 |  |
| (*hsp-16.2(gk249)* Control)/  wildtype Hypoxia | -10.90284 | 6.6112098 | Inf | -1.6491444 | 0.565718 |  |
| (*hsp-16.2(gk249)* Control)/  (*hsp-12.6(ok1077)* Hypoxia) | -35.337753 | 7.9198285 | Inf | -4.4619341 | 0.000118 | *** |
| (*hsp-16.2(gk249)* Control)/  (*hsp-16.2(gk249)* Hypoxia)/ | -7.2089931 | 5.9357633 | Inf | -1.2145014 | 0.829987 |  |
| wildtype Hypoxia/  (*hsp-12.6(ok1077)* Hypoxia) | -24.434913 | 8.4379199 | Inf | -2.8958456 | 0.043843 | * |
| wildtype Hypoxia/  (*hsp-16.2(gk249)* Hypoxia) | 3.6938466 | 6.6112098 | Inf | 0.5587247 | 0.993577 |  |
| (*hsp-12.6(ok1077)* Hypoxia)/  (*hsp-16.2(gk249)* Hypoxia) | 28.12876 | 7.9198285 | Inf | 3.5516880 | 0.005139 | ** |
| **Signif. codes:** 0 ‘***’ 0.001 ‘**’ 0.01 ‘*’ 0.05 ‘.’ 0.1 ‘ ’ 1 | | | | | | |

**Online Resource 22 (Accompanies Fig. 5j)** Statistical analysis for differences in the coefficient of variation of Proteostat fluorescence in the posterior intestine due to genotype and hypoxia treatment. a) Raw data were fitted to a generalized linear model. The R package, ‘emmeans’ was used to obtain estimated marginal means on the response scale (b) and pairwise contrasts (c) with Tukey corrected p-values. Raw data are provided as a Source Data file

**a) Posterior Intestine Coeff. of Variation—Model Statistics**

| **Family:** gaussian (identity) | | | | | | | |
| --- | --- | --- | --- | --- | --- | --- | --- |
| **Formula:** CV ~ Strain * Treatment | | | | | | | |
| **Dispersion:** ~ Strain | | | | | | | |
| **AIC:** -67.8 | **BIC:** -51.9 | | **logLik:** 42.9 | | -**2*log(L):** -85.8 | **df.resid:** 34 | |
| **Dispersion estimate for gaussian family (sigma^2):** 0.00457 | | | | | | | |
| **Conditional model:** | | **Estimate** | | **Std. Error** | **z value** | **Pr(>\|z\|)** | |
| (Intercept) | | 0.90220 | | 0.05150 | 17.518 | < 2E-16 | *** |
| Genotype *hsp-12.6(ok1077)* | | -0.20845 | | 0.05971 | -3.491 | 0.000481 | *** |
| Genotype *hsp-16.2(gk249)* | | -0.13431 | | 0.05732 | -2.343 | 0.019118 | *** |
| Treatment Hypoxia | | -0.32202 | | 0.07284 | -4.421 | 9.81e-06 | *** |
| Genotype *hsp-12.6(ok1077)*: Treatment Hypoxia | | -0.02914 | | 0.08390 | -0.347 | 0.728319 |  |
| Genotype *hsp-16.2(gk249)*: Treatment Hypoxia | | -0.12048 | | 0.08106 | -1.486 | 0.137191 |  |
| **Dispersion model:** | | **Estimate** | | **Std. Error** | **z value** | **Pr(>\|z\|)** |  |
| (Intercept) | | -2.0702 | | 0.2041 | -10.142 | <2e-16 | *** |
| Genotype *hsp-12.6(ok1077)* | | -0.3309 | | 0.2607 | -1.269 | 0.2044 |  |
| Genotype *hsp-16.2(gk249)* | | -0.7164 | | 0.2887 | -2.482 | 0.0131 | * |
| **Signif. codes:** 0 ‘***’ 0.001 ‘**’ 0.01 ‘*’ 0.05 ‘.’ 0.1 ‘ ’ 1 | | | | | | | |

**b) Posterior Intestine Coeff. of Variation—Estimated Model Means**

| **Genotype** | **Treatment** | **emmean** | **SE** | **df** | **asymp.LCL** | **asymp.UCL** |
| --- | --- | --- | --- | --- | --- | --- |
| wildtype | Control | 0.9021981 | 0.051502 | Inf | 0.80125555 | 1.00314072 |
| *hsp-12.6(ok1077)* | Control | 0.6937434 | 0.030204 | Inf | 0.63454348 | 0.75294341 |
| *hsp-16.2(gk249)* | Control | 0.7678865 | 0.025159 | Inf | 0.718574 | 0.81719902 |
| wildtype | Hypoxia | 0.5801775 | 0.051502 | Inf | 0.47923493 | 0.68112011 |
| *hsp-12.6(ok1077)* | Hypoxia | 0.3425808 | 0.028654 | Inf | 0.28641878 | 0.39874282 |
| *hsp-16.2(gk249)* | Hypoxia | 0.3253810 | 0.025159 | Inf | 0.27606857 | 0.37469359 |

**c) Posterior Intestine Coeff. of Variation—Contrasts**

| **contrast** | **estimate** | **SE** | **df** | **t.ratio** | **p.value** |  |
| --- | --- | --- | --- | --- | --- | --- |
| wildtype Control /  (*hsp-12.6(ok1077)* Control*)* | 0.2084546 | 0.0597059 | Inf | 3.4913544 | 0.006391 | ** |
| wildtype Control /  (*hsp-16.2(gk249)* Control*)* | 0.1343116 | 0.0573193 | Inf | 2.3432173 | 0.176843 |  |
| wildtype Control /  wildtype Hypoxia | 0.3220206 | 0.0728352 | Inf | 4.4212221 | 0.000142 | *** |
| wildtype Control /  (*hsp-12.6(ok1077)* Hypoxia) | 0.5596173 | 0.0589370 | Inf | 9.4951776 | 5.04E-14 | *** |
| wildtype Control /  (*hsp-16.2(gk249)* Hypoxia) | 0.5768170 | 0.0573193 | Inf | 10.063222 | 6.73E-14 | *** |
| (*hsp-12.6(ok1077)* Control)/  (*hsp-16.2(gk249)* Control) | -0.0741431 | 0.0393108 | Inf | -1.8860733 | 0.410699 |  |
| (*hsp-12.6(ok1077)* Control)/  wildtype Hypoxia | 0.1135659 | 0.0597059 | Inf | 1.9020866 | 0.400744 |  |
| (*hsp-12.6(ok1077)* Control)/  (*hsp-12.6(ok1077)* Hypoxia)/ | 0.3511626 | 0.0416341 | Inf | 8.4344767 | 5.98E-14 | *** |
| (*hsp-12.6(ok1077)* Control)/  (*hsp-16.2(gk249)* Hypoxia) | 0.3683623 | 0.0393108 | Inf | 9.3705101 | 5.80E-14 | *** |
| (*hsp-16.2(gk249)* Control)/  wildtype Hypoxia | 0.1877089 | 0.0573193 | Inf | 3.2747945 | 0.013511 | * |
| (*hsp-16.2(gk249)* Control)/  (*hsp-12.6(ok1077)* Hypoxia) | 0.4253057 | 0.0381327 | Inf | 11.153287 | 1.87E-14 | *** |
| (*hsp-16.2(gk249)* Control)/  (*hsp-16.2(gk249)* Hypoxia)/ | 0.4425054 | 0.0355814 | Inf | 12.436397 | 0 | *** |
| wildtype Hypoxia/  (*hsp-12.6(ok1077)* Hypoxia) | 0.2375967 | 0.0589370 | Inf | 4.0313673 | 0.000786 | *** |
| wildtype Hypoxia/  (*hsp-16.2(gk249)* Hypoxia) | 0.2547964 | 0.0573193 | Inf | 4.4452104 | 0.000127 | *** |
| (*hsp-12.6(ok1077)* Hypoxia)/  (*hsp-16.2(gk249)* Hypoxia) | 0.0171997 | 0.0381327 | Inf | 0.4510482 | 0.997663 |  |
| **Signif. codes:** 0 ‘***’ 0.001 ‘**’ 0.01 ‘*’ 0.05 ‘.’ 0.1 ‘ ’ 1 | | | | | | |
